# Proteostasis Modelling using Deuterated Water Metabolic Labeling and Data-Independent Acquisition Mass Spectrometry

**DOI:** 10.1101/2025.11.25.690247

**Authors:** Henock M. Deberneh, Daniel J. Wilkinson, Hannah Crossland, Natan Basisty, Kenneth Smith, Philip J. Atherton, Rovshan G. Sadygov

## Abstract

We present the first application of a deuterated water metabolic labeling workflow coupled with data-independent acquisition (DIA) tandem mass spectrometry (MS/MS) for quantifying label enrichment in MS/MS to study protein turnover. The approach automates the turnover rate determination from combined precursor and fragment ions. The truncation of the observed isotope distributions of fragments is overcome by implementing an approach to determining the label enrichment from two mass isotopomers. The high redundancy of fragment ions provides a confident assessment for deuterium enrichment. The DIA approach provides increased proteome coverage and depth for quantifying protein turnover rates compared to the traditional data-dependent (DDA) approach. This novel approach is validated in a murine myotube model of muscle hypertrophy and atrophy through the treatment with insulin-like growth factor 1 (IGF-1) and dexamethasone (Dex), respectively.

## Introduction

Metabolic labeling with deuterated (“heavy” or D_2_O) water followed by liquid chromatography coupled to tandem mass spectrometry (LC-MS-MS/MS) is used to determine individual protein turnover rates in high-throughput^1, 2^. It is used in studies of proteome turnover *in vivo* and in cell cultures^3^. For *in vivo* applications, D_2_O labeling has advantages (compared to labeling with essential amino acids^4–6^ or labeling using a ^15^N diet^7–9^) in cost-effectiveness, ease of use (no diet adaptation is needed), fast equilibration of isotope enrichment^10^, and slow decay of the water pool (permitting expansive measurement periods).

D_2_O-enriched (2-5% in animals, 1-2% in humans, and ∼8% in cell cultures) water is provided as drinking water or a cell medium. The conventional approach is to analyze the labeled samples using high-resolution and mass accuracy mass spectrometers (HRMS) in data-dependent acquisition (DDA) mode. The label incorporation is estimated by using the changes in the isotope profiles of peptides resulting from the deuterium incorporation. Due to incomplete enrichment of the tracer (deuterated water), deuterium incorporation into non-essential amino acids is not complete. This results in the composite spectrum of unlabeled and labeled forms of a peptide in the MS_1_ domain. Bioinformatics techniques have been developed to extract the label enrichments from the isotope profiles^10, 11^. Practical applications have shown that the proteome complexity and semi-stochastic nature of DDA affected the accuracy of turnover rate estimation and proteome coverage^12^. Despite the LC separation and HRMS, contamination of the target isotope profiles by the co-eluting species is common in the samples from mammalian proteomes^13^. The techniques that use two mass isotopomers^14, 15^ (instead of the complete isotope profiles) have increased the depth of proteome (for turnover studies) coverage by about 40%. The spectral accuracy (accurate reproduction of the isotope profile) of the peptides in MS1 is also affected by the space-charging, saturation, and limited dynamic range. In addition, the semi-stochastic nature of DDA leads to the well-documented missing data problem^16^. While the match between runs approach to transfer the peptide matches between experiments mitigates the problem^17^, there are errors associated with the procedure^18^. One possible strategy that addresses the semi-stochastic nature of DDA and co-eluting profiles of peptides is the data-independent acquisition (DIA)^19^. In the DIA approach, all precursors in a predefined mass range are fragmented. Peptide spectrum matching (PSM) is performed using rich fragment data information in the tandem mass spectra (MS/MS)^19–21^. DIA has been shown to be consistent for peptide identification and quantification.

Here, we report on the use of MS1 and MS/MS spectra in DIA to determine protein turnover rates of deuterated water labeled samples, **Figure 1**. Proof-of-concept with novel methodologies such as these requires a level of biological validation at the cellular level. To address the efficacy of this novel DIA analysis, we focused on skeletal muscle as an example of a highly adaptive tissue that can exhibit rapid alterations in phenotype (i.e., through changes in protein turnover in response to external stimuli). For instance, anabolic or catabolic effectors, where tissue protein mass may be rapidly gained^22^, or lost^23^, respectively. Such responses were mimicked within a C2C12 murine myotube model treated with the insulin-like growth factor 1 (IGF-1) to promote hypertrophy through simulation of protein synthesis, and the glucocorticoid dexamethasone (DEX) to promote atrophy through the suppression of protein synthesis.

**Figure 1.**
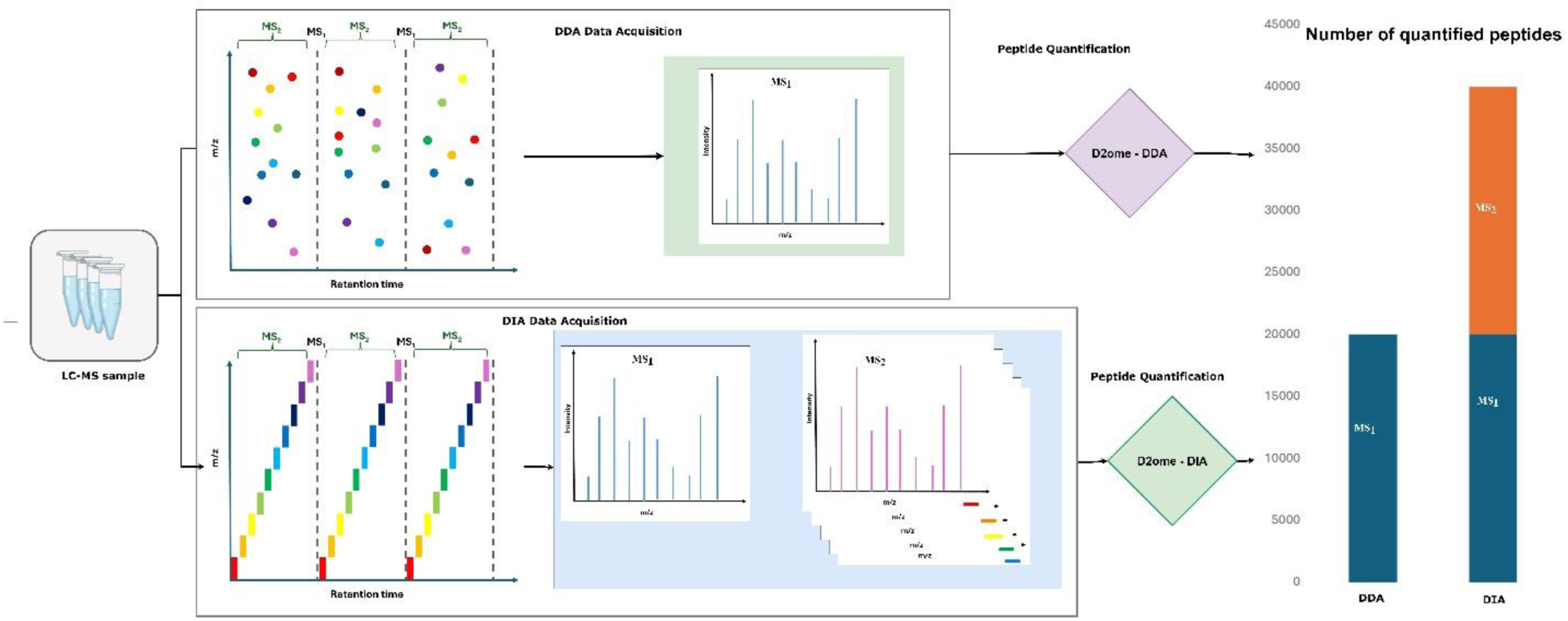
DIA acquisition increases the coverage and depth of proteome in protein turnover studies using deuterated water metabolic labeling. DIA-data acquisition produces fragment ions enriched in deuterium from deuterated water. The isotope distributions of the fragment ions are used to compute the label incorporation. The quantification from the fragment ions increases the depth and coverage of the cellular proteome.

## Results

### Myotube Morphology of Culture Cells

To confirm the effectiveness of the two treatments on inducing cell anabolism (IGF-1) and inhibition of cell growth (DEX) within the fully differentiated C2C12 myotubes, light microscopy images were used to measure the changes in myotube width (surrogate measure of cell size) due to the treatments at 24h and 48h. There were significant decreases in myotube width compared to control cells at both 24h (21.5±0.6 vs. 16.2±0.5; *P*<0.001 vs. Control; **Supplementary Figure 1**) and 48h (22.2±0.6 vs. 18.5±0.5; *P*<0.001 vs. Control; **Supplementary Figure 1**) with DEX treatment, and a contrasting significant increase in myotube width compared to control cells at 24h (21.5±0.6 vs. 25.0±0.8; *P*<0.001 vs. Control; **Supplementary Figure 1**) and 48h (22.2±0.6 vs. 25.7±0.8; *P*<0.001 vs. Control; **Supplementary Figure 1**) with IGF-1 treatment.

### The comparison of proteome coverage for protein turnover in DIA and DDA

The performance of a proposed DIA approach was evaluated by directly comparing its quantification output with the conventional DDA approach across three different time-course LC-MS experiments using a C2C12 murine myotube model of hypertrophy and atrophy, **Figure 2**. These experiments involved treatments with IGF-1, DEX, and a control group. The time-course experiments were conducted over five distinct labeling durations: 0, 4, 8, 24, and 48 hours, with up to six technical replicates for each duration. Each experiment was performed twice, once in DDA mode and once in DIA mode. The raw output files from the LC-MS experiments were converted to the mzML^24, 25^ format using the MSConvert tool of proteowizard^26^ and subsequently used as input for the database search engines. To identify proteins and peptides in the DDA experiments, we used the MASCOT^27^ database search engine, while the DIA experiments were analyzed using the DIA-NN^20^ database search engine for DIA spectra. To quantify deuterium enrichment of peptides in the DDA experiments, we used d2ome software (d2ome-DDA), whereas the proposed tool for DIA experiments, d2ome-DIA, was utilized for analyzing DIA data.

**Figure 2.**
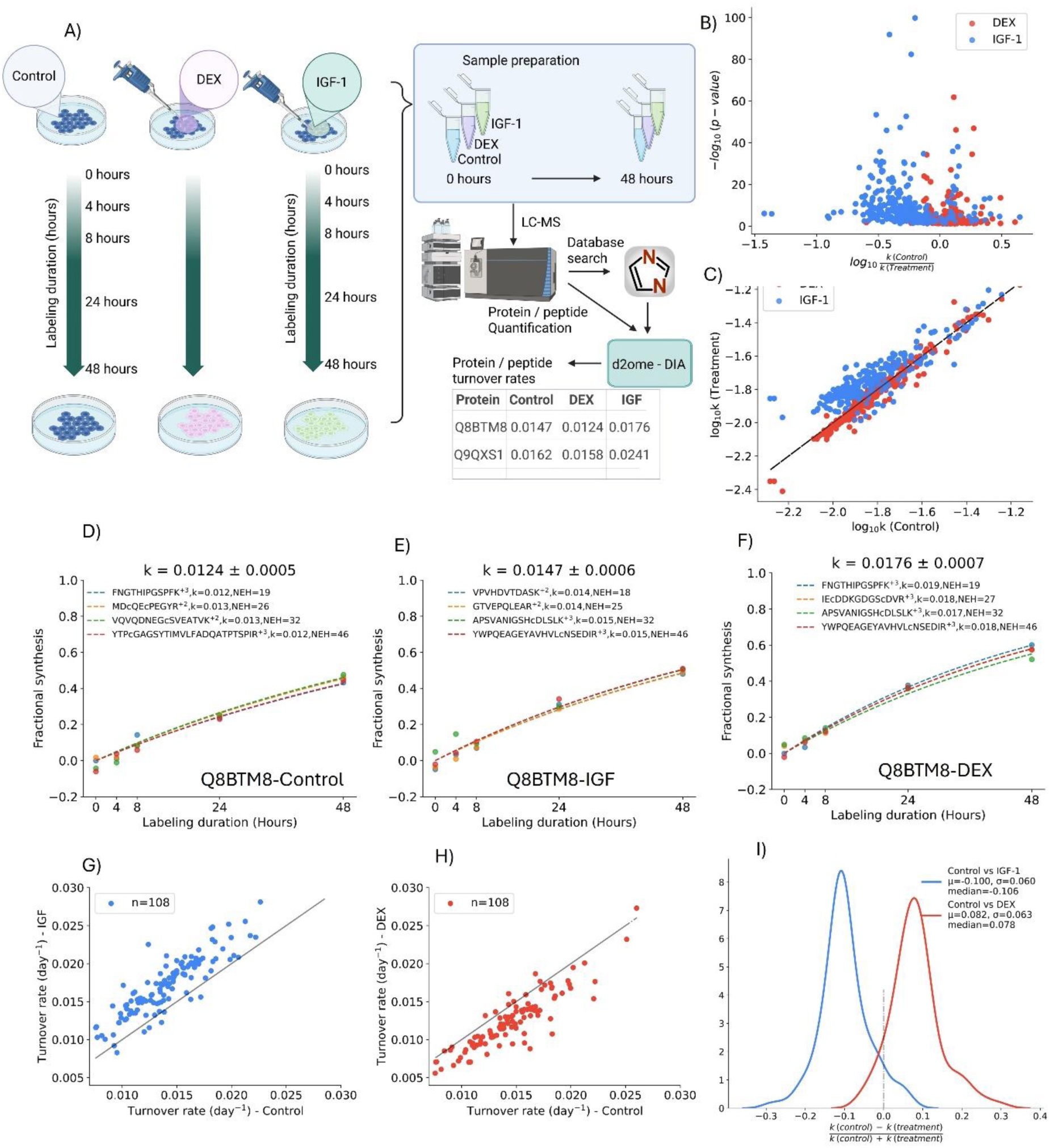
The workflow of D_2_O labeling and comparison of the turnover rates of proteins in insulin-like growth factor 1 (IGF-1) and dexamethasone (DEX) treated cells with those in the control cells. Shown are turnover rates for 268 proteins that were common in the three conditions. **A**) Experimental approach, **B**) Volcano plots of turnover rates of common proteins, **C**) Scatter plot of the turnover rates, and **D)** time course data and turnover rates of protein Fibronectin type-III domain-containing protein 3A (Q8BTM3), control, **E**) IGF-1-treatement, **F**) DEX treatment; **G**) scatter plots of the control and IGF-1 treated **G**) and control and **H**) DEX-treated samples for peptides of Q8BTM3; **I**) the density plots of the DEX-treated and IGF-1 treated peptide relative turnover rates for Q8BTM3.

The new approach, d2ome-DIA, consistently quantified a significantly higher number of proteins compared to the DDA approach across all three experimental conditions, **Figure 3**. In the control experiments, d2ome-DIA quantified 6,986 proteins (**Supplementary Data 1**), while d2ome-DDA quantified only 2,682 (**Supplementary Data 2**). A similar trend was observed in the DEX experiment, where d2ome-DIA quantified 7021 proteins (**Supplementary Data 3**) compared to 2,752 by d2ome-DDA (**Supplementary Data 4**). Furthermore, for the IGF-treatment, d2ome-DIA quantified 6216 proteins (**Supplementary Data 5)**, whereas d2ome-DDA quantified only 2,690 (**Supplementary Data 6**). **Supplementary Figure 2** illustrates the common and unique protein quantifications, highlighting that a substantial number of proteins are uniquely quantified by DIA in each condition. For example, in the control experiment, DIA exclusively identified 4,591 proteins, while 2,679 were shared by both methods.

**Figure 3.**
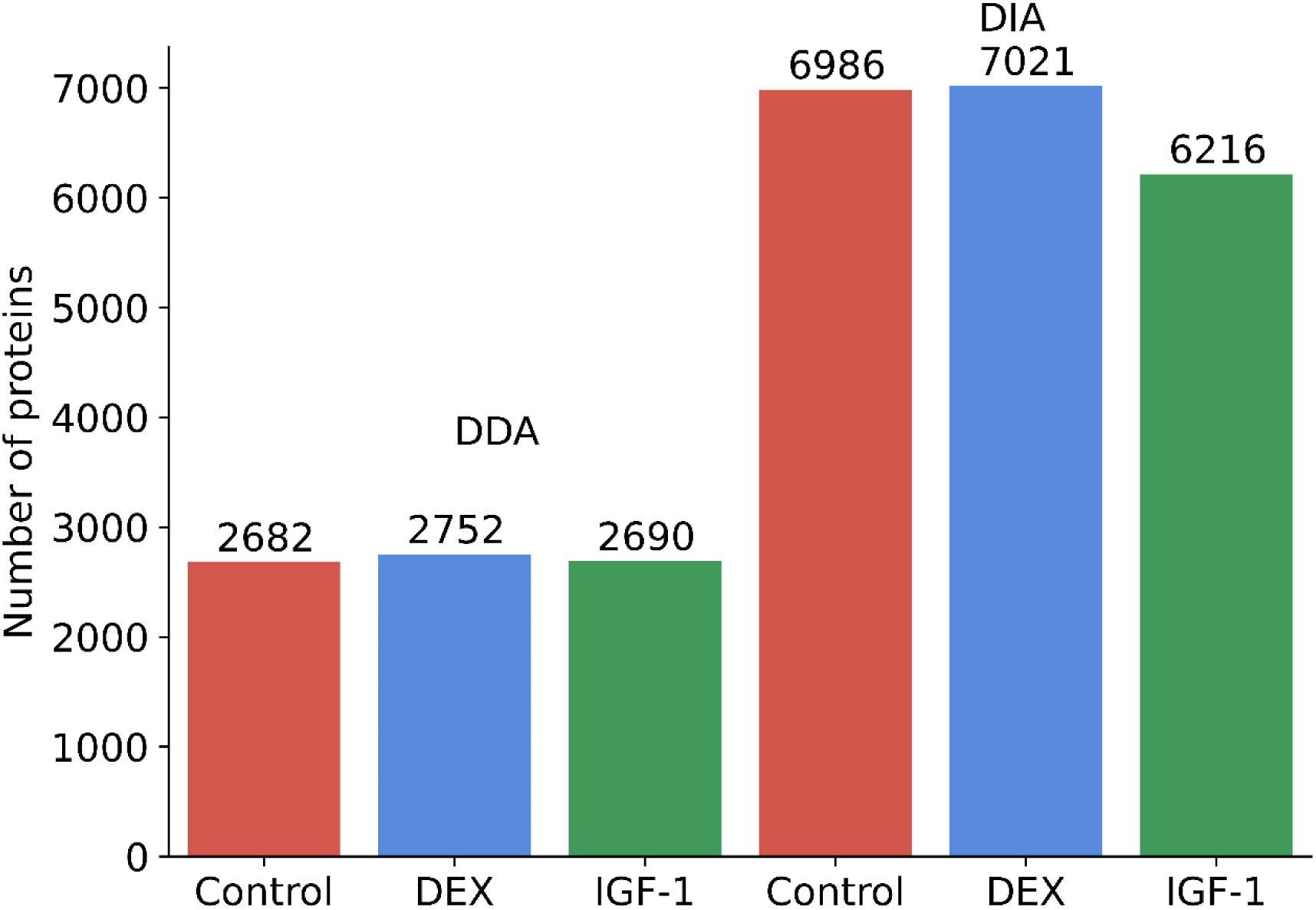
The numbers of proteins identified in data-independent acquisition mode (DIA) is more than twice as many as in data-dependent (DDA) acquisition methods used in this work. Most proteins quantified in DDA were also quantified in DIA, whilst in the DIA, we uniquely quantified ∼1.7 times more proteins. DEX – dexamethasone, IGF-1 insulin-like growth factor 1.

d2ome-DIA consistently outperformed the DDA method in the number of quantified peptides (**Supplementary Figures 3**). For example, in control experiments, the DDA approach quantified 25,003 peptides from 2,682 proteins, while d2ome-DIA quantified 73,773 peptides (**Supplementary Figure 3 A**) from 6,986 proteins. Only distinct peptides are counted in comparing the performances of the DDA and DIA approaches.

The Venn diagrams of the quantified peptides for the control experiment (**Supplementary Figure 3 B - D**) illustrate a significant overlap between the DDA and DIA approaches but also highlight a substantial number of peptides unique to DIA. The two-way Venn diagram (**Supplementary Figure 3 B**) shows that DIA uniquely identified 51,393 peptides, with 22,380 peptides common to both DDA and DIA. The three-way Venn diagram (**Supplementary Figure 3 D**) for DIA-MS1, DIA-MS/MS, and DDA shows that 21,878 peptides were common to all three methods.

This pattern is consistent across the DEX and IGF treatment groups. **Supplementary Figure 4** shows that d2ome-DIA identified 74,406 peptides, while d2ome-DDA identified 25,573. The corresponding Venn diagrams (**Supplementary Figure 4 B - D**) confirm the above trend, showing a significant overlap of 22,797 peptides but a much larger number of unique peptides identified by DIA, totaling 51,609. This trend persists in the IGF-1 condition (**Supplementary Figure 5**), where DDA identified 24,710 peptides and DIA identified 59,832 peptides. The Venn diagrams show that 21,774 peptides are common to both DDA and DIA, with DIA identifying 38,058 unique peptides.

We note that the number of proteins and peptides quantified in the DDA method in this study is comparable with the corresponding numbers in samples from cell cultures^3^ and *in vivo* experiments,^13^ which used D_2_O labeling. For example, 18,392 peptides from 2,995 proteins were quantified in a recent study of protein turnover in human induced pluripotent stem cells (hiPSC)^3^ using D_2_O labeling. A different study^28^ by Cambridge and colleagues using SILAC^29^ labeling (a conventional approach for labeling proteins with heavy essential amino acids in cell cultures) monitored the depletion of existing proteins following the cycloheximide inhibition of new protein synthesis and quantified the turnover rates of 3528 proteins (quantified by at least two distinct peptides) in C2C12 cells. The turnover rates of proteins were determined from two labeling points: 0 and 24 hours of labeling with SILAC. The median of the turnover rates of all proteins in the study was 0.016 day^-1^. From the DDA method, we obtained a similar median (in the control sample) for the turnover rates of all quantified proteins, 0.0156 day^-1^. The median of the turnover rates of all proteins quantified in the DIA method was slightly higher, 0.02 day^-1^. As the results show, overall, both acquisition methods from our work agreed well with the cellular proteome turnover rate from the previous study.

### The Comparison of Peptide Turnover Rates (linear regression)

This section provides a detailed analysis of the peptide turnover rate comparison between DDA and DIA using only MS1 (referred to as MS1), and MS/MS (also referred to as MS2) approaches for the datasets obtained from control sample. To quantify deuterium enrichment, we used the linear regression model (see the Methods section). The results for DEX and IGF-1 treated samples, as well as the isotope averaging method, are similar and provided in the **Supplementary Information** file.

To quantify the goodness-of-fit between the experimental time course data and theoretical fit, the coefficient of determination (R^2^) is used. Higher R^2^ values indicate a close fit. It is customary to require that a peptide be quantified in at least four points of labeling and have R^2^ ≥ 0.8. We used these criteria in our evaluation of the turnover rate estimations.

The pie charts in **Figure 4** illustrate the distribution of R^2^ values of peptides across the DDA (MS1 spectra), and MS1 and MS/MS spectra in DIA for the control sample. The corresponding pie charts for DEX and IGF-1 treated samples are shown in **Supplementary Figures 6 A** and **B**, respectively. In the control sample (**Figure 4**), the percentage of peptides with a R^2^ ≥ 0.95 was the highest for the MS/MS spectra of DIA method and reached 49.8%, followed by DDA at 44.9% and MS1 of DIA at 45.1%. While the percentages of highly accurate peptide quantifications are comparable, the absolute numbers are drastically different. Thus, quantification using MS/MS spectra in DIA resulted in more than threefold increase in the number of quantified peptides with high goodness-of-fit (R^2^ ≥ 0.95) than in DDA (34578 vs. 11283). The trend is consistent across the other two conditions (**Supplementary Figures 6 A** and **B**) and indicates that the estimation of peptide turnover rates using MS/MS spectra generally yields a vastly higher number of peptides with very high R^2^ values.

**Figure 4.**
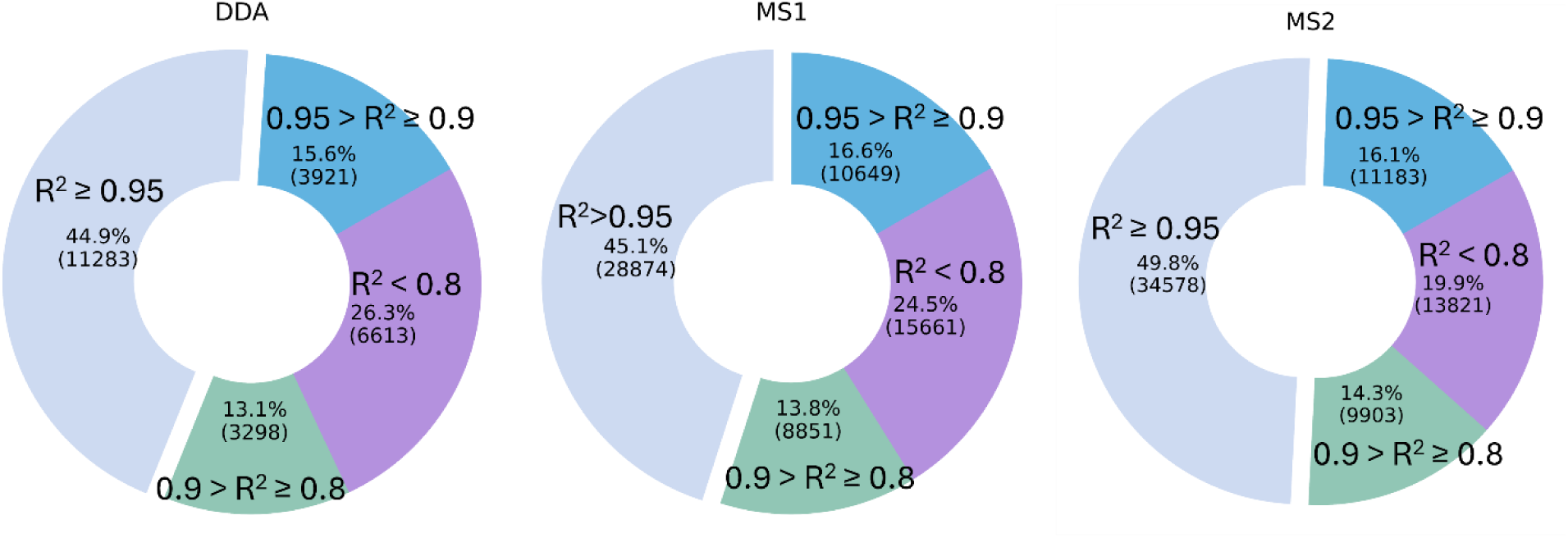
The numbers and percentages of peptides for several intervals of the coefficient of determination (R^2^) between the theoretical fit and experimental time course of the label enrichment for the control sample from data-dependent acquisition (DDA) and full mass spectra (MS1) and tandem mass spectra (MS/MS) of the data-independent acquisition (DIA). The number of peptides quantified with R^2^ > 0.95 from their fragment ions in MS/MS (DIA approach) was more than three times higher than that in DDA.

The scatter plots in **Supplementary Figures 7 A-C** compare the natural logarithm of turnover rates (ln(k)) between the DDA and MS1 of DIA methods for control, DEX-treated and IGF1-treated samples. The peptides were filtered such that R² ≥ 0.9 (n=7834 for control sample). The Spearman correlation was 0.89. Similar trends are observed in the DEX-treated and IGF-treated samples (**Supplementary Figures 7 B** and **C,** respectively), where Spearman correlations were 0.90 (n=7283) for the DEX-treated sample, and 0.87 (n=7569) for IGF-treated sample. The density plots of the relative differences (computed as described above) between turnover rates are shown in **Supplementary Figures 8 A-C**. The median of the differences was about 0.02 for all samples, and the SDs were also similar and approximately equal to 0.22.

The comparison of turnover rates of peptides computed from DDA and MS2 of DIA exhibited more differences, **Supplementary Figures 9 A-C** (the scatter plots of logs) and **Supplementary Figures 10 A-C** (density plots of the relative differences). The medians and SDs from the distributions of differences were larger than those from MS1, 0.12 (median) and 0.51 (SD).

### Uncovering New Biological Insights Unseen with DDA

With a larger number of proteins and peptides able to be identified and quantified using DIA compared with DDA in this cell model, the increased proteome coverage could provide deeper insight into potential mechanisms driving phenotypic change in the cell model, that would not be possible with DDA. Comparing DIA and DDA from the control only rates of turnover, protein accession numbers which were common to both DDA and DIA modes, and those unique to only DIA mode were extracted and mapped to their functional ontologies using clusterProfiler^30^. Of the biological processes mapped to these proteins, 29.6% were uniquely identified to the DIA mode proteins (**Figure 5**). These functions unique to DIA mode were mainly associated with the regulation of RNA metabolism and ribosomal biogenesis in the cell (RNA splicing, RNA catabolic processes, rRNA metabolic processes, mRNA catabolic processes, ribosomal biogenesis; **Figure 6**), key processes involved in the regulation of the synthesis of new proteins within the cell. Whereas functions associated with proteins common to both DDA and DIA modes related primarily to control of energetic processes in the cell required for cellular integrity and assembly (energy derivation by oxidation of organic compounds, ATP metabolic processes, ribonucleotide metabolic processes; **Figure 7**). These data highlight the increased depth of knowledge afforded by DIA, which could lead to novel discoveries when applied to other models of health and disease.

**Figure 5.**
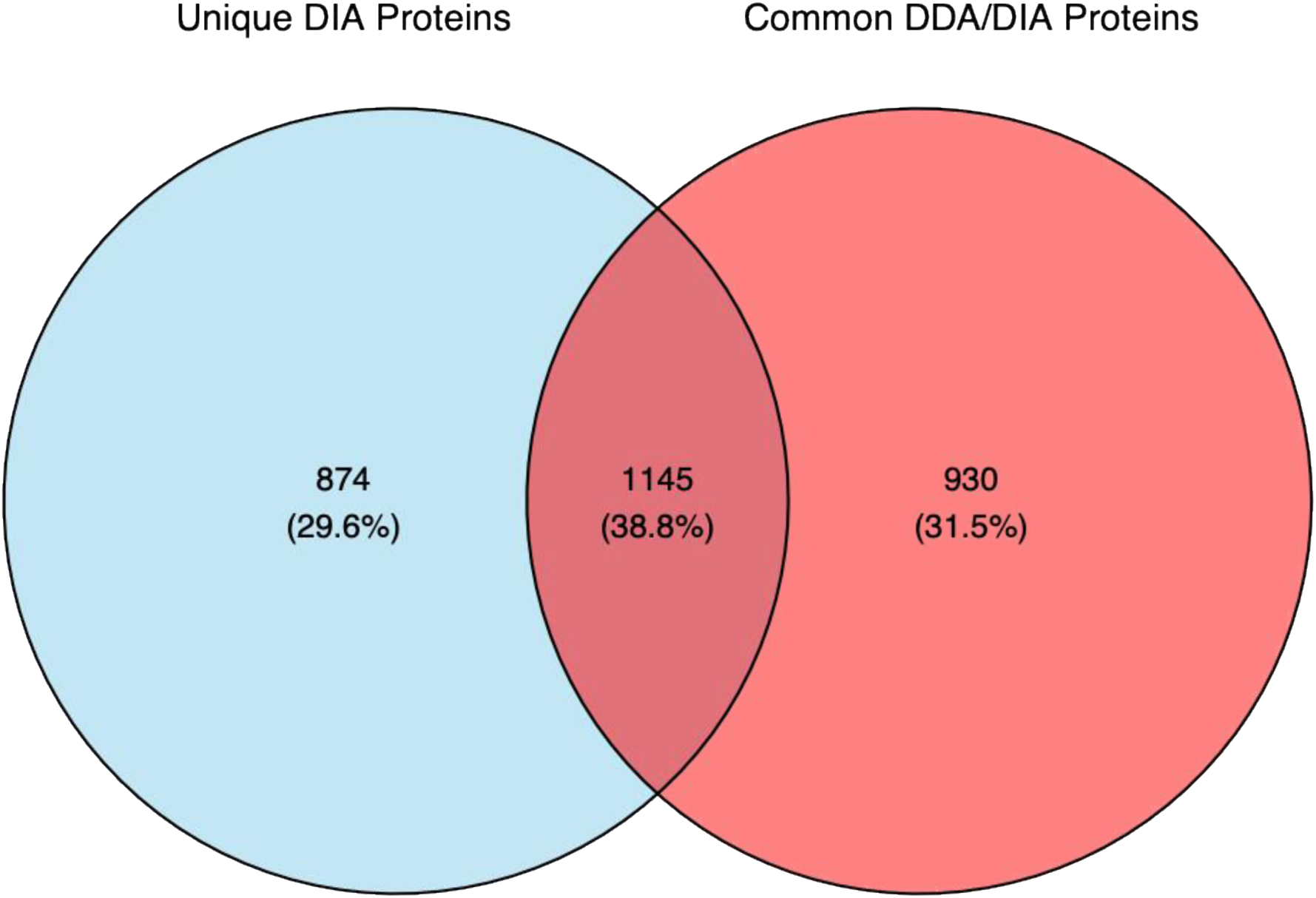
Venn Diagram of overlap between associated functional enrichment for unique and common proteins amongst data-independent (DIA) and data-dependent (DDA) acquisition modes. The left circle (blue) represents the proteins that are unique to pathways generated from DIA-identified proteins. The right circle (red) represents the proteins that are common to the pathways generated by proteins from DDA and DIA. Of the proteins, which were in common pathways for DDA and DIA, 38.8% were also in the pathways that were unique to DIA only. There was no pathway which was unique to DDA-identified proteins only.

**Figure 6.**
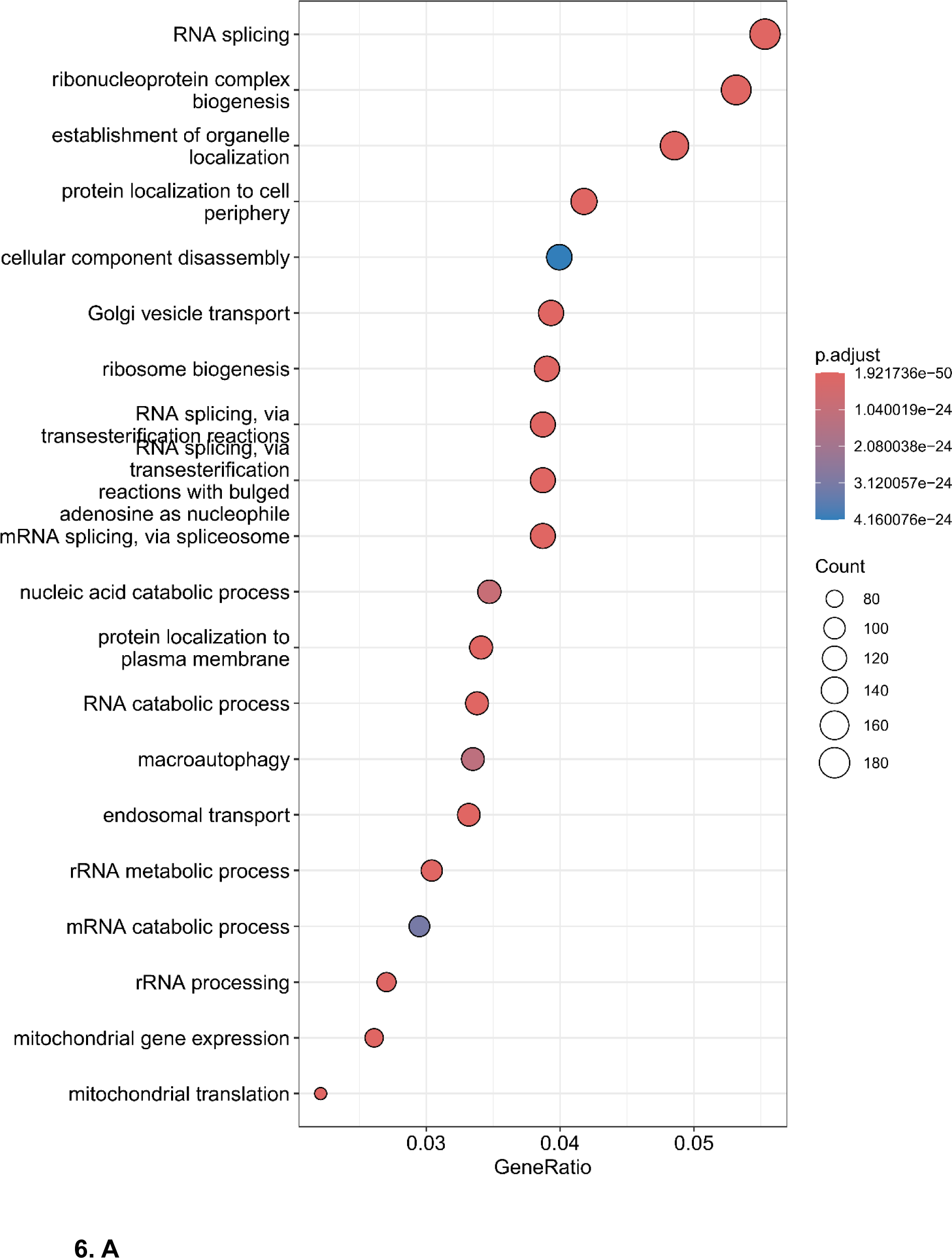
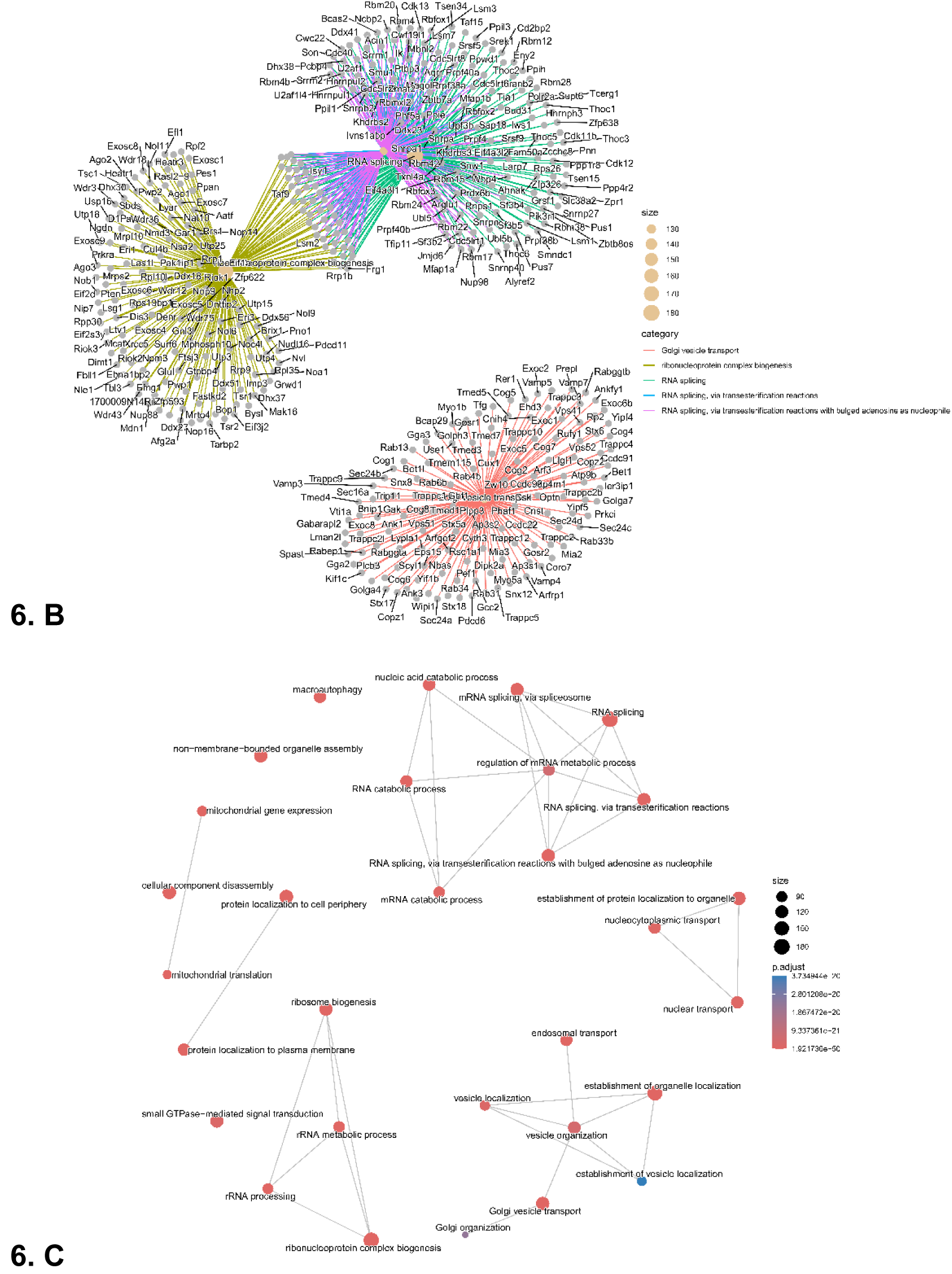
Biological functions associated to proteins quantified only in data-independent acquisition (DIA) mode. A) Dotplot highlighting the top 20 most highly enriched GO terms for proteins common to either mode. B) cnetplot (a function of clusterProfiler) depicting linkages between individual genes and Gene Ontology (GO) enriched terms. C) Enrichment map of networks of interconnecting and overlapping functional gene terms.

**Figure 7.**
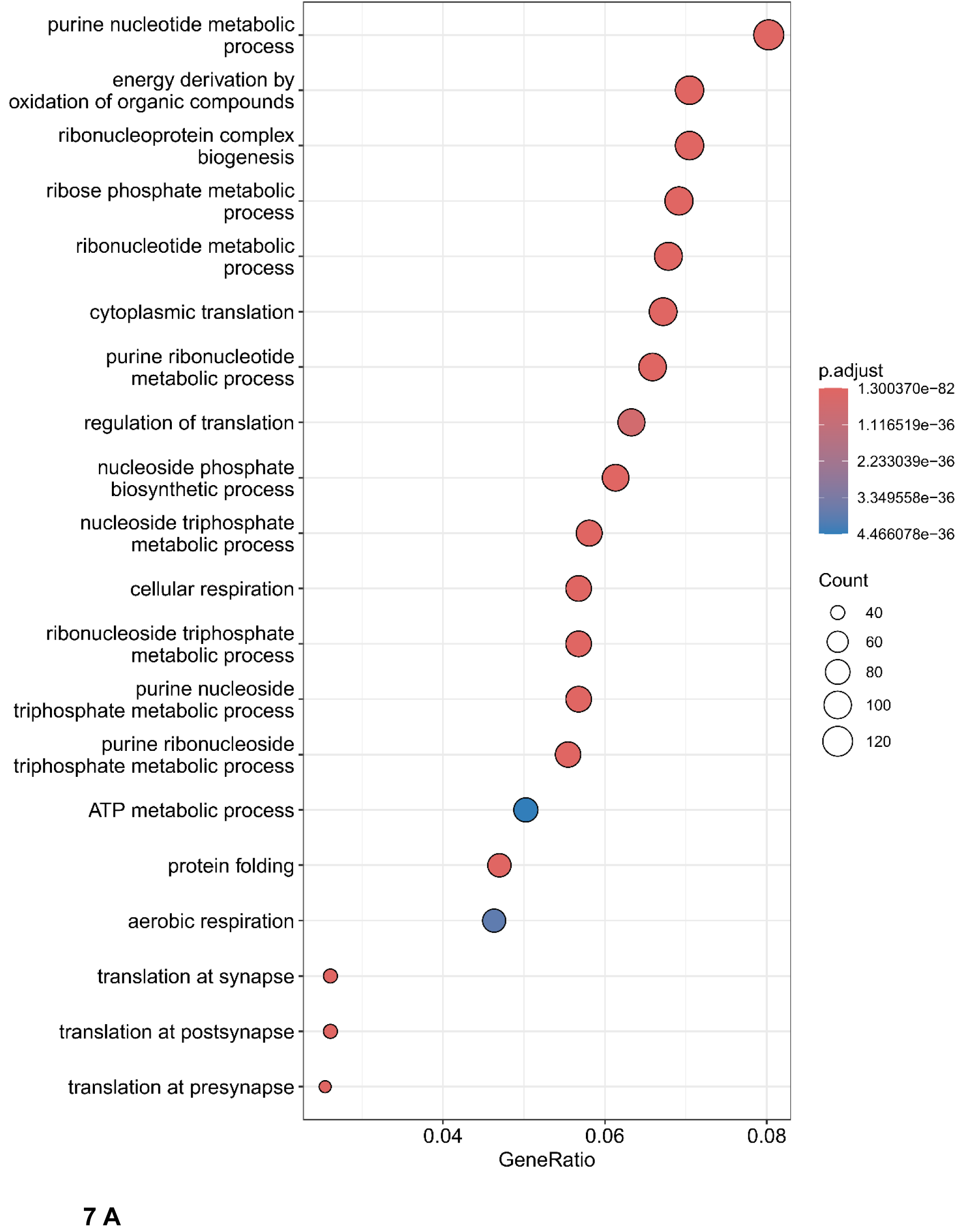
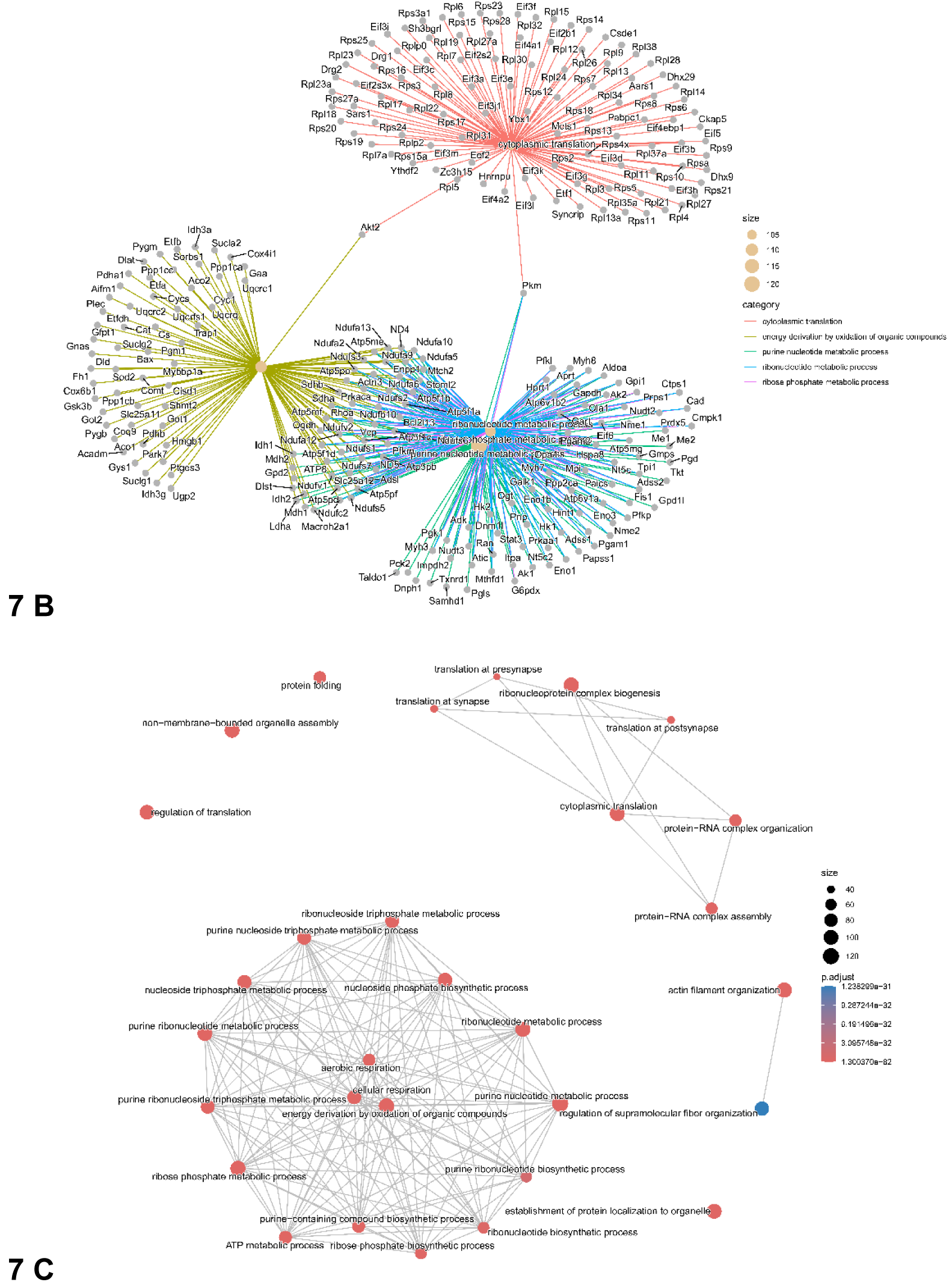
Biological functions associated with proteins common to both data-dependent (DDA) and data-independent (DIA) acquisition modes. A) Dotplot highlighting the top 20 most highly enriched Gene Ontology (GO) terms for proteins unique to DIA mode. B) cnet plot depicting linkages between individual genes and GO enriched terms. C) Enrichment map of networks of interconnecting and overlapping functional gene terms.

### Biological Function and Gene Ontologies of Significantly Changing Protein FSR in DIA mode

For individual proteins found to have significantly changing FSRs due to treatment with either IGF-1 or DEX, we sought broader context by investigating the functional roles of these proteins within the biology of the muscle cell, i.e., mapping the proteins to their gene ontologies and key functions. Following treatment with IGF-1 the vast majority of significantly changing proteins increased in FSR (**Figure 8A**), consistent with an established anabolic effector (the same pattern of change was also observed in DDA mode, however with a lower number of proteins, see **Supplementary Figures 11-15**). Functional enrichment analyses found that these proteins were key to the regulation of ribosomal function (ribonucleotide metabolic process, ribonucleoprotein complex biogenesis) and translational regulation (cytoplasmic translation, regulation of translation) of new proteins (**Figure 9**). Networks also highlighted this, with many ribosomal proteins and eukaryotic initiation and elongation factors identified as significantly changing (**Figure 9**). With DEX treatment, whilst a greater number of proteins decreased in FSR, there were several proteins increasing in FSR as well (**Figure 8B** volcano plots). Proteins significantly increasing in FSR due to DEX treatment were primarily associated with muscle cell structure and integrity (actin filament organization, muscle contraction, actin-filament based movement; see **Figure 10**), while the proteins with significantly decreasing FSR due to DEX treatment were associated with the regulation of mRNA and translation (cytoplasmic translation, RNA splicing) in addition to the regulation of key cellular energetics (NAD and NADH metabolic processes, glycolysis; see **Figure 11**).

**Figure 8.**
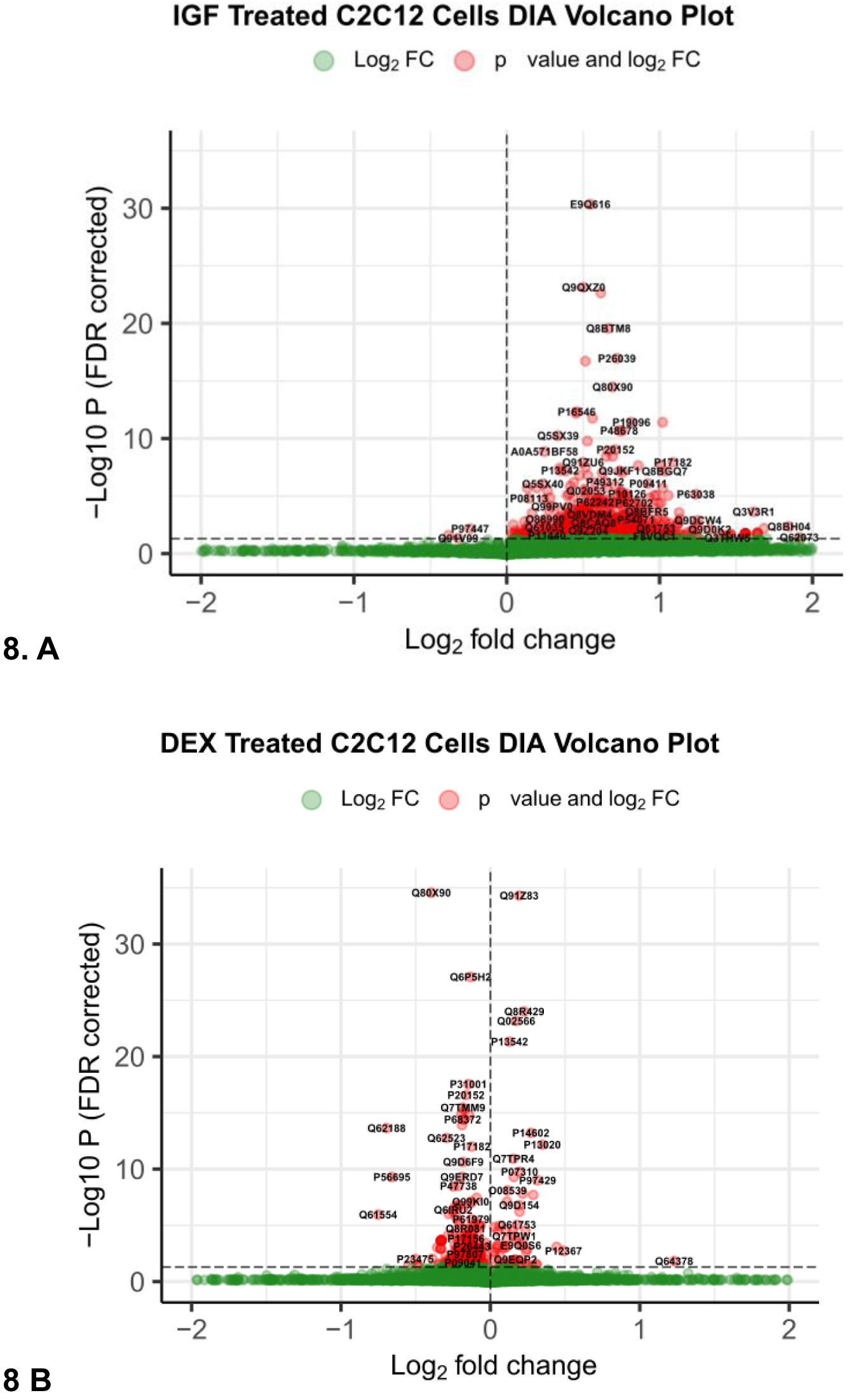
Volcano plots illustrate significantly changing proteins identified and quantified in data-independent (DIA) acquisition mode following treatment with **A**) insulin-like growth factor 1 (IGF-1)-treated sample, and **B**) dexamethasone (DEX) – treated sample.

**Figure 9.**
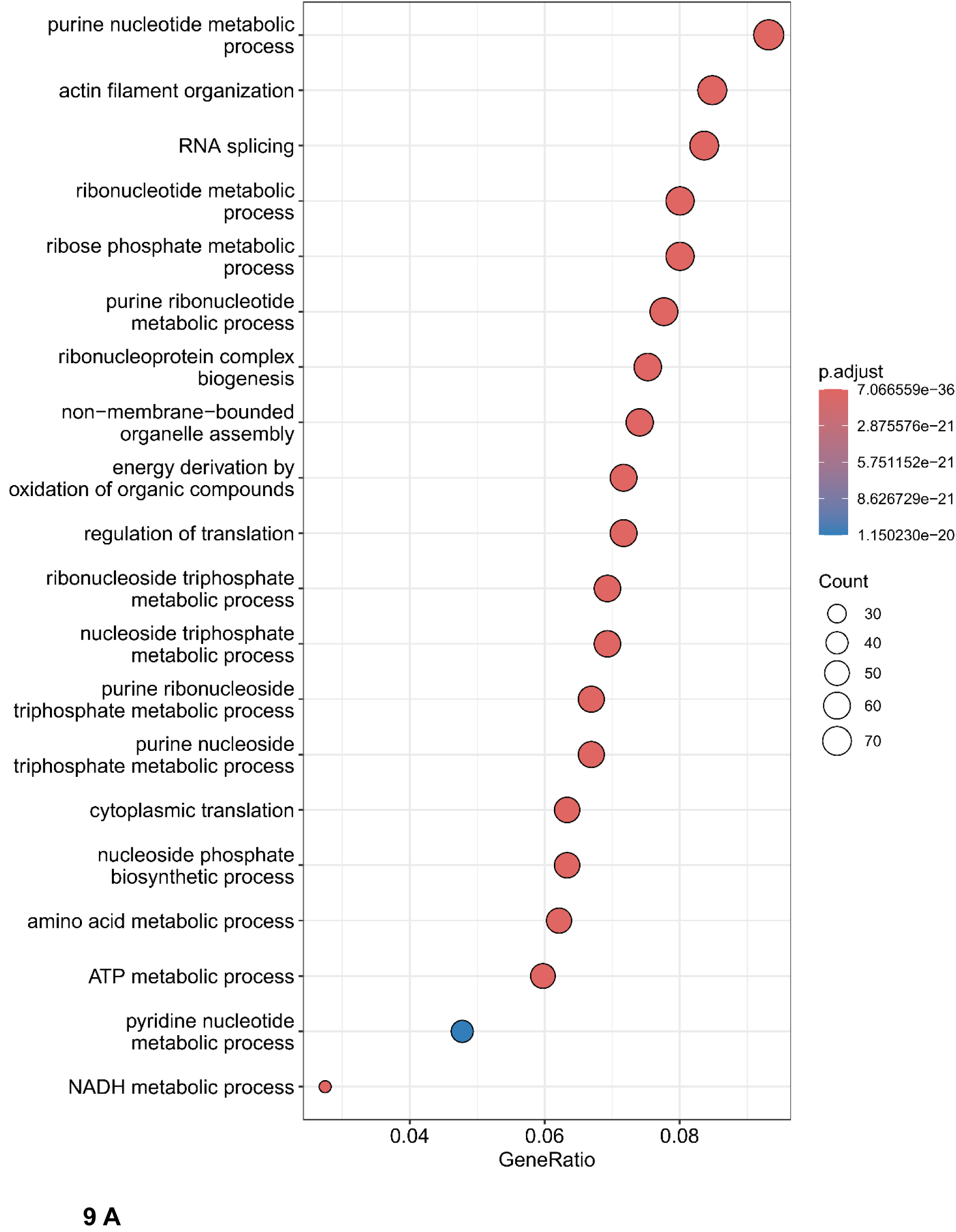
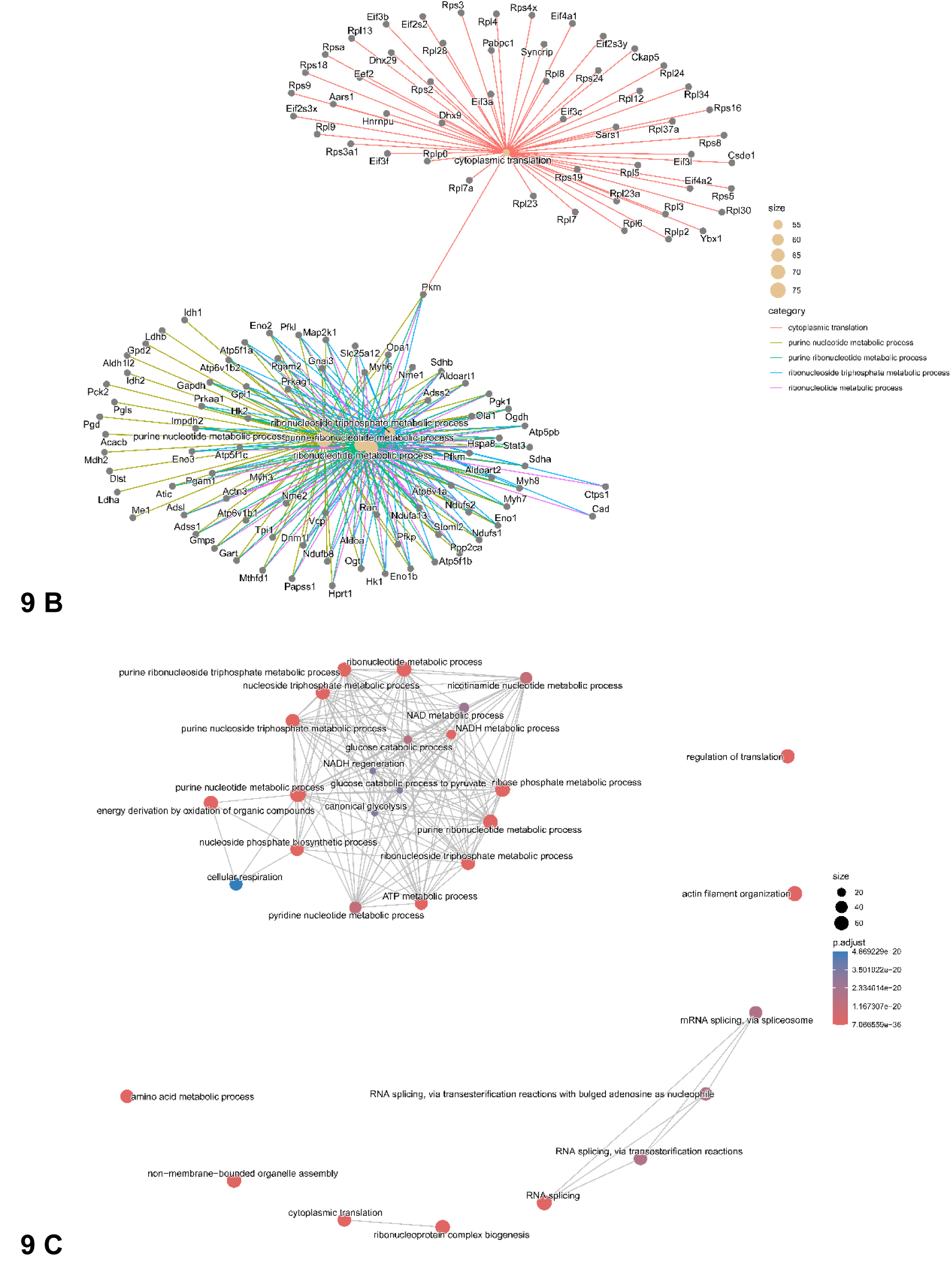
Functional enrichment analyses of insulin-like growth factor 1 (IGF-1) treated C2C12 cells for significantly changing proteins. A) Dotplot highlighting the top 20 most highly enriched Gene Ontology (GO) terms for significantly changing proteins. B) cnet plot depicting linkages between individual genes and GO enriched terms. C) Enrichment map of networks of interconnecting and overlapping functional gene terms.

**Figure 10.**
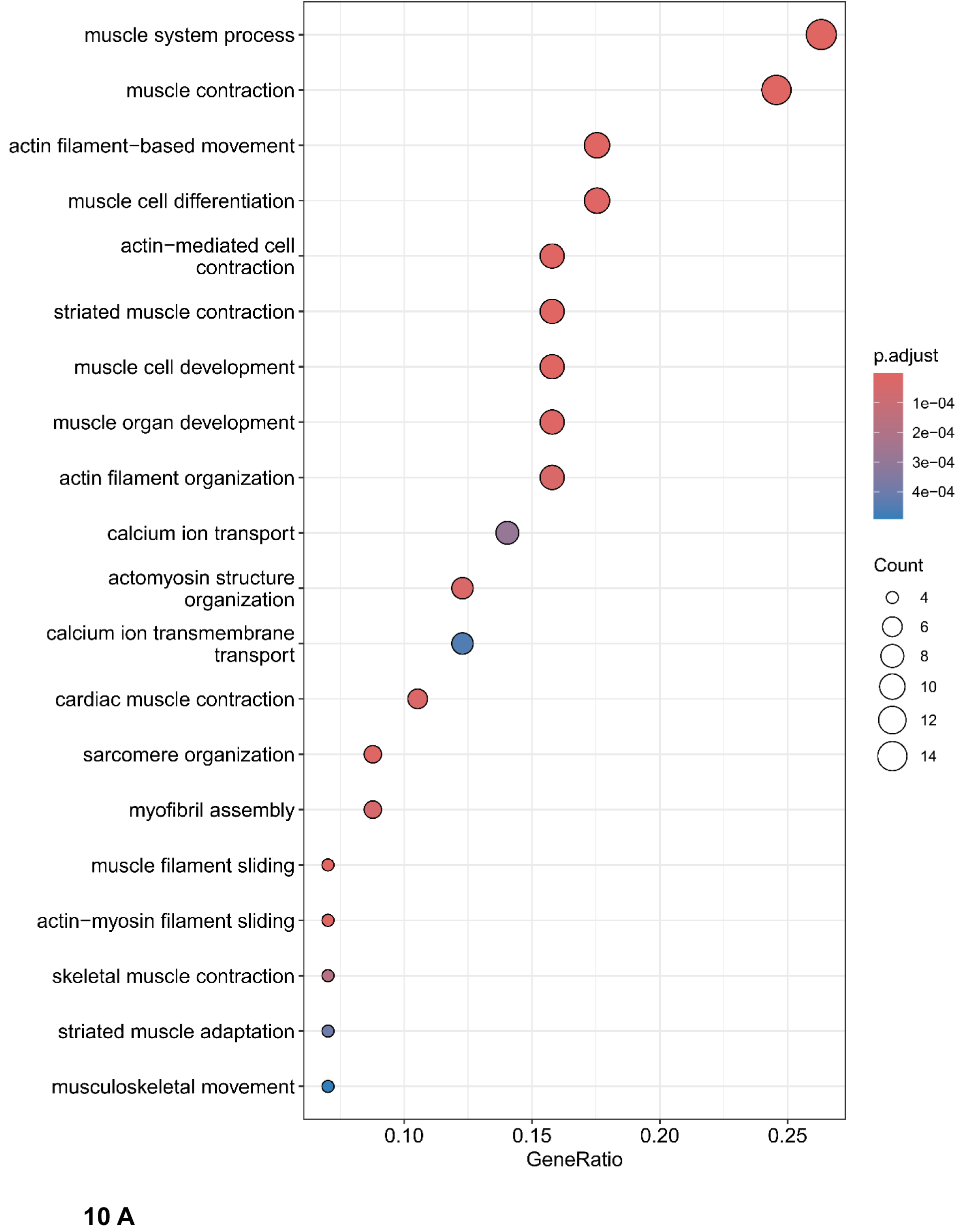
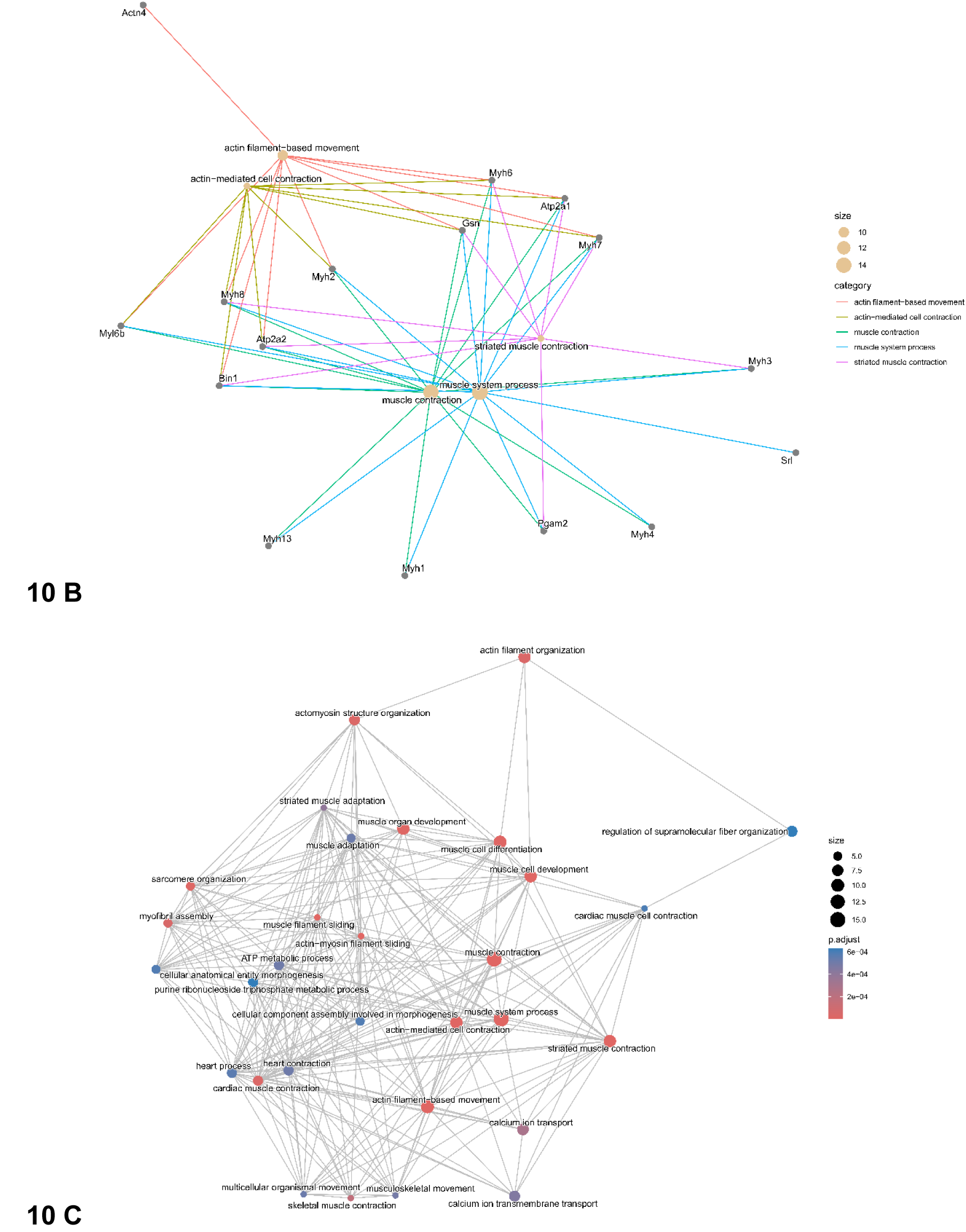
Functional enrichment analyses of proteins significantly increase in fractional synthesis rate (FSR) in response to dexamethasone (DEX) treatment. A) Dotplot highlighting the top 20 most highly enriched GO terms for significantly changing proteins. B) cnet plot depicting linkages between individual genes and GO enriched terms. C) Enrichment map of networks of interconnecting and overlapping functional gene terms.

**Figure 11.**
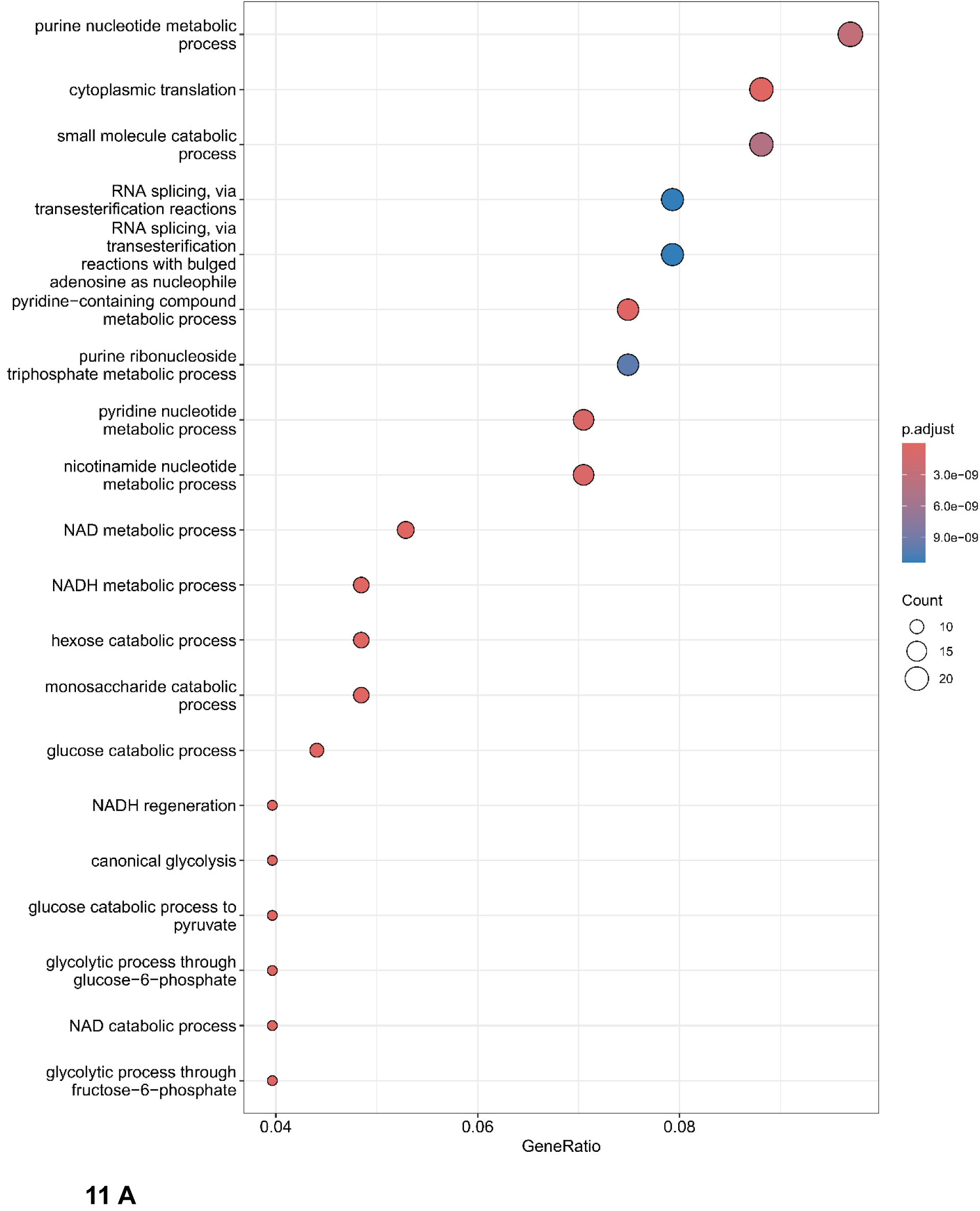
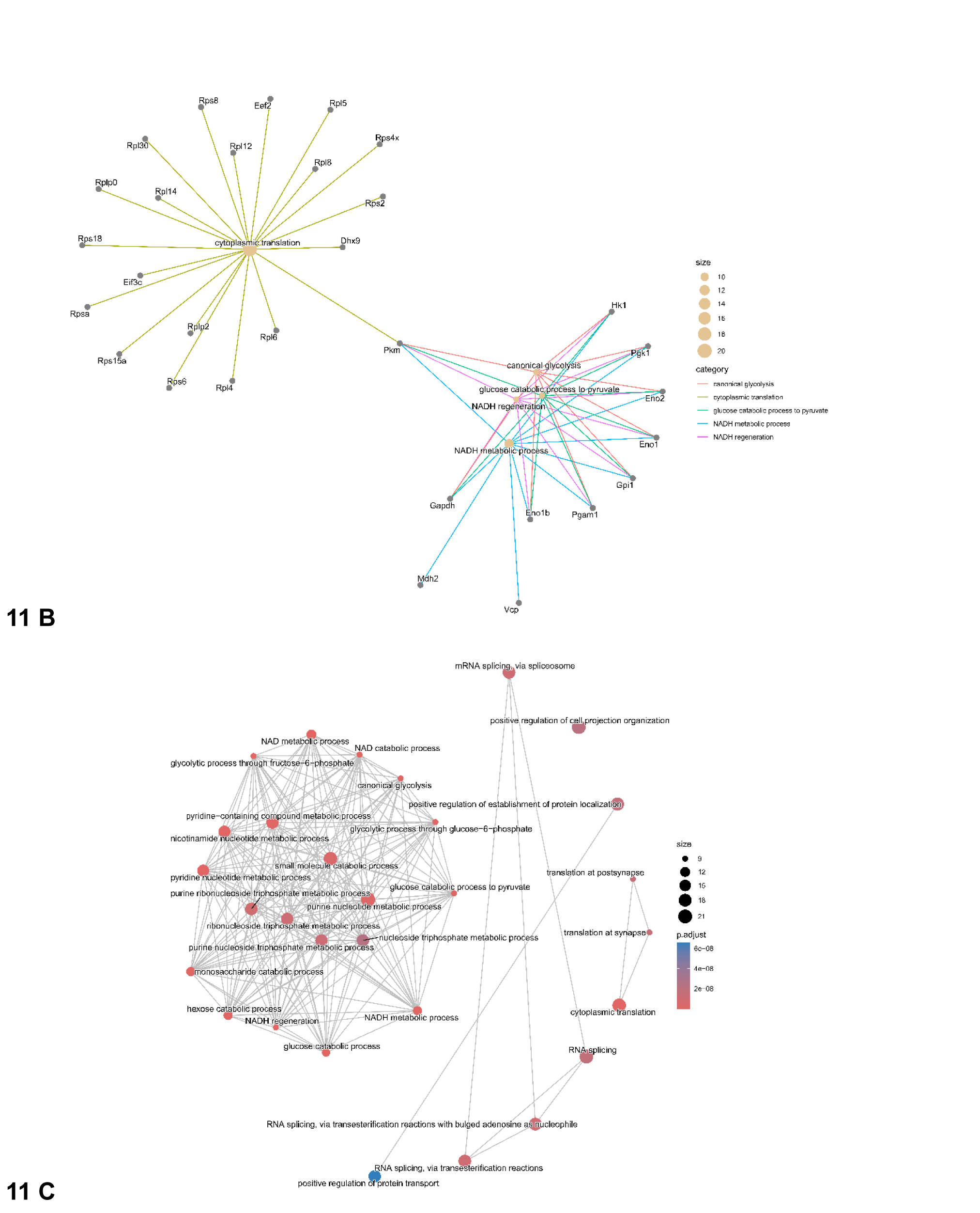
Functional enrichment analyses of proteins significantly decreasing in fractional synthesis rate (FSR) in response to DEX treatment. **A**) Dotplot highlighting the top 20 most highly enriched GO terms for significantly changing proteins. **B**) cnet plot depicting linkages between individual genes and GO enriched terms. **C**) Enrichment map of networks of interconnecting and overlapping functional gene terms.

## Discussion

In this work, we demonstrate how to combine MS1 and MS/MS data in DIA to compute protein turnover rates of deuterated water labeled samples, **Figure 1**. To the best of our knowledge, the work is the first to apply the DIA and MS/MS quantification for estimating protein turnover. A recent study^3^ used DIA for peptide/protein identifications from deuterated water labeled samples. However, no quantification of label enrichment from MS/MS spectra was reported. In our study, the label quantification uses both MS1 and MS/MS spectra. We note that in DIA MS/MS spectra, like in DDA MS/MS spectra, the isotope distributions of fragment ions are often truncated and include only the monoisotope and the first heavy mass isotopomer. Therefore, an approach that can quantify the label enrichment from two mass isotopomers is critical for estimating the labeling using fragment ions. We have previously developed an analytical approach to determine the label enrichment from two mass isotopomers^14^. Here, we use this technique to determine the deuterium enrichment from two mass isotopomers of a fragment ion (also referred as a product ion). The combined use of MS1 and MS/MS increased the proteome coverage by more than two times (number of quantified proteins).

To assess the efficacy of DIA, we focused on skeletal muscle as an example of a highly adaptive tissue that can exhibit rapid alterations in a phenotype. In terms of internal validation, applying anabolic or catabolic stimuli, where tissue protein mass may be rapidly gained^22^, or lost^23^, provides such a tractable model. Clinically, muscle atrophy occurs in myriad clinical conditions, such as malnutrition, intensive care, and acute/chronic disease, such as cancers, organ failures, and infectious diseases^31^. Given how muscle vitality can influence clinical outcomes^32^ there is significant research into skeletal muscle proteostasis in relation to the mechanistic underpinnings of both positive and negative influencers of muscle mass - reflecting our application herein. Moreover, from a cell biology perspective, a benefit of in vitro approaches is the ability to study the direct effects of anabolic/catabolic paradigms strictly in muscle cells, without the influence of knock-on systemic effects, tissue crosstalk, or influences of non-muscle cells (e.g., the murine C2C12 cell line^33, 34^). Taking the opportunity to study established and clinically relevant drivers of hypertrophy (insulin-like growth factor 1 (IGF-1^35–37^) and atrophy corticosteroids (DEX^38^), that induce opposing changes in muscle mass and protein pool sizes, we addressed their relative influences on muscle proteostasis to provide the first dynamic OMIC readouts in these scenarios.

### The effects of IGF-1 and DEX treatments on protein turnover rates

IGF-1 induces muscle hypertrophy by binding to its cognate receptor (IGF-1R) and activating a canonical PI3K-Akt-mTOR signaling pathway^35^. As anticipated, treatment of C2C12 cells with IGF-1 induced an increase in myotube diameter reflecting protein accretion and muscle hypertrophy, as previously reported^37^. Reflecting this, expansion of the total proteome mass is a causal feature of muscle hypertrophy, which herein can be explained by the gross upregulation of individual protein synthesis uncountered by reciprocal downregulation i.e., 918 proteins upregulated and 65 downregulated. Such a bulk upregulation of muscle proteins being synthesized in response to a trophic stimulus (e.g., IGF-1^37^, nutrition, exercise^39^) is also observed with the use of traditional amino acid stable isotope tracing when extracting proteins from a tissue mass and measuring the collective synthesis of whole-crude protein fractions, such as sarcoplasmic, myofibrillar, collagen and mitochondrial pools^40, 41^. That structural molecule activity was among the largest functional enrichment (**Figure 9**) substantiates a bulking up of contractile/cytoskeletal components to complement increased cellular mass^42^. Reciprocal to this, molecular function analyses highlighted a marked upregulation of protein synthesis associated with transcription (initiation factors), mRNA binding, and ribosomal assembly, consistent with activation of IGF-1 canonical pathways^35^ and stimulation of mTORc1. In addition to the activation of muscle protein synthesis (MPS), reflecting the rate/efficiency of mRNA translation, is the capacity for MPS in the context of ribosomal content, or ‘capacity’ for MPS. Ribosomal protein content has been shown to increase in response to muscle growth stimuli^43^ and is functionally necessary to sustain increased rates of MPS. Indeed, mTORC1 controls the translation of terminal oligopyrimidine (TOP) mRNAs, which code for ribosomal proteins and several initiation and elongation factors, by acting on La-related protein 1 (LARP1), a key repressor of TOP mRNA translation^44^. IGF-1 is therefore causing growth via upregulating the capacity for protein synthesis.

Treatment of muscle cells with DEX led to a characteristic atrophy of myotubes, as previously reported^38^ via its characteristic binding to intracellular glucocorticoid receptors inducing a transcriptional programme causal in muscle atrophy, suppressed MPS and transcriptional regulation of proteolytic factors (myostatin, and the E3 ligases ‘atrogenes’, MuRF-1 and MAFBx^45^). In contrast to IGF-1, DEX mostly downregulated the synthesis of proteins (64 up and 246 down), likely reflecting a shrinking proteome mass and the observed atrophic response. That actin and cytoskeletal binding functional clusters were regulated likely reflects sarcomere remodeling and disassembly^46^ as characteristic features of muscle atrophy induced by prevailing protein degradation. In terms of other major functional aspects, altered RNA binding (top 3 downregulated ontologies) and 3’ UTR binding was evident. As such, the posttranscriptional regulation of gene expression by RNA-binding proteins (RBPs) and microRNAs (miRNA) involved in the turnover and translation of mRNA could be involved in the induction of muscle atrophy, as has been shown in aging muscle tissue^47^. In summary, application of these novel methodologies yielded restructuring of the dynamic proteome that could explain regulatory features of protein accretion with IGF-1 and protein loss with DEX. This work paves the way for deep dynamic phenotyping (identification of pathways, key proteins etc.) regulating cellular proteostasis in a multi-modal and tissue agnostic fashion.

The comparison of the turnover rate calculations with a previous study^28^

Cambridge and colleagues^28^ quantified turnover rates of 3563 proteins from C1C12 cell culture using SILAC labeling. There were 2121 and 3206 common proteins between the Cambridge et al. work^28^ and proteins quantified in DDA and DIA methods, respectively, in our study. 121 protein entries from Cambridge et al. study are not currently distributed in the murine protein sequence database by uniport (most of these entries were retired or will be retired by 2026). When we required quantification by at least two distinct peptides in our results from DDA and DIA methods, there were 1901 (DDA) and 3069 (DIA) common proteins between our and Cambridge et al. results. It is important to point out that in SILAC labeling experiments quantified proteins were filtered to have at least two distinct peptides. The experiments were done in two labeling times points: 0 and 24 hours. In our studies, we required proteins to be quantified in at least four time points of labeling. We, too, filtered quantified proteins to have at least two distinct peptides. In addition, we required that proteins to have turnover rates less than 0.07 h^-1^, as in the SILAC experiments the fastest turnover rate was 0.066 h^-1^. The above shown numbers of common proteins (between DIA/DDA and SILAC experiments) were obtained after the application of these filters. The density plots of the relative difference between turnover rates from our study (control sample) and SILAC labeling are shown in **Supplementary Figure 16**. The relative difference was computed as the difference between turnover rates from the two studies divided by their sum. **Supplementary Figure 16 A** shows the results obtained using the linear regression (see the Methods section for linear regression and isotope averaging methods to estimate label enrichment from fragment ions), and **Supplementary Figure 16 B** shows the results from isotope averaging. The comparisons to SILAC results from both approaches to label enrichment estimations are similar. As shown in **Supplementary Figure 16 A**, the median of the relative differences of protein turnover rates was 0.04 and the standard deviation (SD) was 0.23 for quantification from DIA. The D_2_O labeling using DIA for sample analysis shows robust agreement with SILAC data. The corresponding data from the DDA approach showed similar agreement: the median of the relative differences was 0.07 and the SD was 0.22. At the modes of the relative difference distributions, the turnover rates in both methods of our study differed from those in the SILAC labeling experiments by approximately 4% to 7%. Spearman correlation between the turnover rates from our study and SILAC labeling was 0.7 for both DDA and DIA turnover rates. The p-values of the Spearman correlations were less than 10^-258^. As mentioned above, DIA quantified 50% more common (with the SILAC experiments) proteins than DDA did.

## Methods

### Data modeling

#### Turnover rate estimation from precursor ions

The turnover rate of proteins and peptides from precursor ions (in MS1) is determined using exponential decay modeling of the time course of the depletion of the relative abundance (RA) of the monoisotope^48^. Traditionally, the monoisotopic RA is quantified for several labeling durations (datapoints for at least four labeling durations are used) from a complete isotope profile. The experimental time-course data are regressed against the exponential decay function:

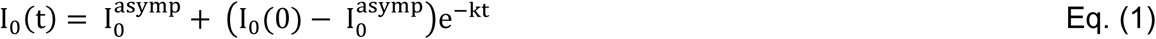

where k is the turnover rate, I_0_^asymp^ is the monoisotopic RA at the plateau of labeling (computed theoretical), I_0_(0) is the monoisotopic RA of natural peptide (with the natural deuterium abundance), and t is the labeling duration. I_0_^asymp^ is computed using the body water enrichment in deuterium (p_W_), the number of exchangeable hydrogens/labeling sites (N_EH_), and I_0_(0)^49^:

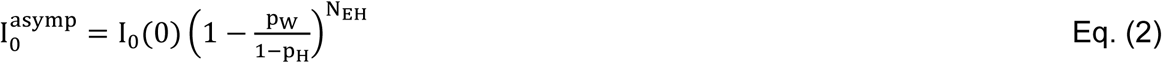

In Eq. (2), p_H_ is the natural abundance of ^2^H. The turnover rate, k, is obtained from the non-linear least-squares regression^50^ of the experimental time course data on the decay function in Eq. (1). The quality of the estimated turnover rate is evaluated based on the experimental to theoretical goodness of fit criteria such as coefficient of determination (R^2^), root mean squared error (RMSE), and Pearson correlation coefficient^51^. The variance of the turnover rate is estimated using the delta method^13^.

### The properties of fragment ions of metabolically labeled peptides

In DDA, which uses only a limited window of isolation of precursor ions, only a partial isotope profile of a peptide is chosen for fragmentation. For example, the often-used 1.4 Th window centered on the monoisotope would fully include the monoisotope itself and the first heavy mass isotopomer in the fragmentation of a +2 charged peptide. Therefore, the isotope distributions of resulting fragment ions in DDA do not correctly reflect the isotope content of the precursor peptide^52^. The problem is especially acute for the deuterium-labeled samples, as the labeling leads to an increase in the average mass of a peptide, often resulting in a change in the mode of peptides’ isotope distribution. This problem is not present in DIA, as a wide m/z window is used to select precursors. It could still be present if the selected fragmentation window does not fully cover the isotope profile of a precursor (for a precursor at the edges of the fragmentation window). The selection of the overlapping isolation windows will resolve the issue.

Isotope distributions of fragment ions are often truncated and contain only the monoisotope and the first heavy mass isotopomer (**Supplementary Figure 17**). Therefore, we used our recently developed approach to estimate the label enrichment (excess enrichment in deuterium, p_X_(t)) using two mass isotopomers only. p_X_(t) is determined from the raw abundances of the first two mass isotopomers in the isotope distribution:

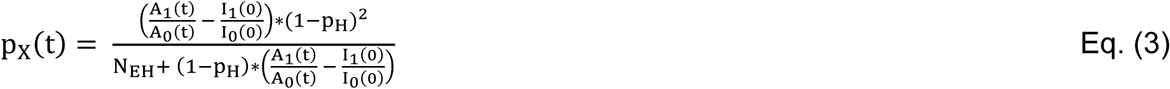

The time course of the p_X_(t) is related to the turnover rate using the following equation:

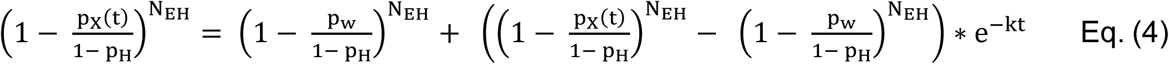

Thus, using Eq. (3), we estimate the excess enrichment in the deuterium via the abundances of the first two mass isotopomers of a fragment ion. Then from the time course of the excess deuterium enrichment of a fragment ion, Eq. (4), we obtain the turnover rate.

The comparison of label enrichments from the fragment ions and the precursor ion can be obtained using the level of deuteration, N_EH_*p_X_(t). It is expected that the deuteration of fragment ions will be on a line drawn from the center to the deuteration value of the precursor ion. This property of the deuteration is reflected in **Supplementary Figure 18**, where we show the deuteration values of fragment ions and the precursor ion of the peptide, “EGFFELIPQDLIK^+2^”. As seen from the figure, the relative deuteration levels of fragments lie on a line, which connects the center (of the coordinate system) and the deuteration level of the precursor ion. The figure is an example demonstrating the validity of our approach for extracting the deuterium enrichment level from the fragment ions of a peptide. The linearity property is also used in the linear regression analysis (see below) in which we extract the enrichment level of a peptide from its multiple fragments.

In **Supplementary Figure 19**, we show the label enrichment estimations from MS1 (the precursor ion) and MS/MS (multiple fragment ions) spectra for several labeling durations and six replicates at each labeling duration for peptide, LDEAEQLALK²⁺ of murine protein Myosin-8. As seen from the figure, the lines connecting median of enrichment levels as calculated from MS1 and MS2 closely match. As expected, the modeling of the turnover rates from these data produces similar results.

The tandem mass spectrum of a peptide contains multiple fragment ions. Each fragment ion represents a part of the peptide sequence. The availability of many fragment ions for quantification provides the advantage of applying statistical approaches to determine turnover rates. However, it also poses a challenge in identifying ions that exhibit correct isotope profiles reflecting label incorporation. Often, fragment ions have lower intensity than precursor ions, and their isotope profiles contain two or at most three mass isotopomers. Only 3.34% of fragment ions had a complete isotope profile from the control dataset. Consequently, applying a method similar to that used for precursor ions (complete isotope profile) to estimate turnover from fragment ions becomes challenging. To address this challenge, we have implemented two computational approaches in this study.

### Isotope averaging

We developed a technique to determine the excess deuterium enrichment by convolving fragment ion intensities. For example, for two fragment ions, we compute the averaged ratio of the first heavy mass isotopomer to the monoisotope, 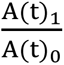, as:

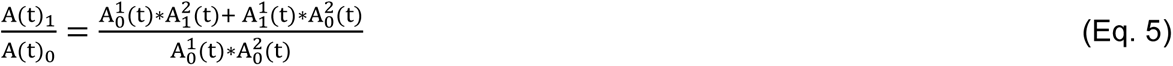

where A(t)_i_ represents the intensity of the i^th^ mass isotopomer of the convolved signal, 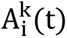 represents the intensity of the i^th^ mass isotopomer of the k^th^ fragment ion. The N_EH_ corresponding to the averaged fragment mass isotopomers is the sum of the N_EH_ values of all fragment ions used in the quantification. The excess deuterium enrichment was computed using Eq. (5).

The intensities of the mass isotopomers of the fragment ions were quantified from MS/MS spectra using the start and end elution times of the precursor ion reported in the database search report file of DIA-NN^20^. For each MS/MS scan within the elution window of a peptide, the isotope profile of up to four fragment ions was quantified, and the convolved ratio, 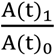, was calculated. We then compute the p_x_(t) values for each scan and take the median value as the representative label incorporation value for the given labeling duration. Eq. (3) was used to determine the p_x_ (t) values from 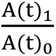. This process is repeated for each labeling duration to determine the time course p_x_(t) values. The turnover rate from fragment ions is determined using non-linear regression on the time course label incorporation values, p_x_(t), determined from MS/MS spectra using Eq. (4).

### The excess deuterium enrichment estimation from fragment ions: linear regression

This approach utilizes linear regression to determine the label incorporation values, p_x_(t), for a given labeling duration. Like the previous method, we start with extracting the isotope profile of each fragment ion in every MS/MS scan within the elution window of a peptide. For the fragment ion, we compute the difference between the ratios of the first heavy and monoisotopic ions in labeled and unlabeled samples. We store this information for every fragment ion. Next, we use the linear relationship between the ratio difference and the N_EH_ value as shown below:

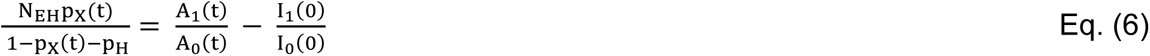

The regression of the experimental values for fragment ions on the right-hand side of Eq. (6) over the N_EH_ values of different fragments of a peptide produce the expression, 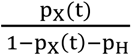, from which we determine the deuterium enrichment, p_X_(t).

### Software Inputs and outputs/DIA Quantification workflow

D2ome-DIA is a stand-alone software tool designed to compute turnover rates for proteins and peptides from heavy water-labeled data-independent LC-MS experiments. The primary inputs for the software are LC-MS spectral files in the “mzML” format^26^ and database search results in comma-separated (.csv) file format. D2ome-DIA is compatible with DIA-NN database search outputs and accepts results from any search engine, provided they are preprocessed as specified in **Supplementary Table 1**. The database search results must contain the following information: peptide sequence, start elution time, end elution time, precursor mass, and a list of the top four fragment ions along with their corresponding quality scores. In the output, the code provides the ability to view the chromatographic elution profiles (for manual evaluation) of the precursor ion and fragment ions of a peptide. The details of the workflow are described in the Supplementary Note. **Supplementary Figure 20** illustrates the graphical user interface (GUI). **Supplementary Figure 21** shows the GUI of the experimental time course and theoretical fit generated for a protein by d2ome. **Supplementary Figure 22** demonstrates the ability of the d2ome-DIA to visualize the chromatographic elution profiles of quantified peptides at each labeling duration from every replicate mass spectrometry run. **Supplementary Figure 23** shows the elution profiles of the precursor and fragment ions of the peptide LDEAEQLALK²⁺ (MYH8_MOUSE).

### C2C12 Cell Culture and MS Sample Preparation

Murine C2C12 cells (passage 6, ECASS, Salisbury, UK) were maintained in Dulbecco’s Modified Eagles Medium (DMEM, Thermo Fisher Scientific) containing 10% (v/v) fetal bovine serum (FBS), 1% antibiotic-antimycotic solution, with 4mM l-glutamine (All from Sigma-Aldrich, UK). Myoblasts were seeded onto collagen-coated (Type I from rat tail; Sigma-Aldrich, UK) six-well multi-dishes (Nunclon™ Delta; Thermo Fisher Scientific) and when confluency reached ∼90%, differentiation was induced by changing to DMEM containing 2% (v/v) horse serum, 1% (v/v) antibiotic-antimycotic solution and 4 mM l-glutamine (All Sigma-Aldrich, UK). The media was changed every 48 h and experiments were performed at days 4 post differentiation.

At the start of the experiment, media was replaced with DMEM containing 10 Atom Percent Deuterium Oxide (D_2_O) stable isotope tracer (ISOTEC, Merck Life Sciences, UK) to measure protein turnover (as previously described^37^), in addition to the following treatments: Untreated controls (Ctrl: 4 plates of n = 6 per plate technical replicates), IGF-1 50ng/ml final concentration (IGF: 4 plates of n = 6 per plate technical replicates), Dexamethasone 2.5uM final concentration (DEX: 4 plates of n = 6 per plate technical replicates). One plate per treatment was harvested at 4h, 8h, 24h, and 48h following initial treatment into cell lysis buffer (Easy Pep MS Sample Prep Kit, Thermo Fisher Scientific, Hemel Hempstead, UK). An additional plate with no stable isotope tracer added was also harvested to act as the 0 h for all treatments and provide baseline natural abundance measures for protein turnover calculations.

Following harvest, samples were normalized to 100ug of protein, processed and digested to their constituent peptides using the Easy Pep Mini MS Sample Prep Kit (Thermo Fisher Scientific, Hemel Hempstead, UK) according to the manufacturer’s instructions. Following sample preparation, samples were evaporated to dryness under nitrogen and reconstituted in 100ul of 95% Water and 5% Acetonitrile with 0.1% Formic Acid (All Optima LC-MS Grade, Fisher Scientific, Loughborough, UK) ready for MS analyses.

### MS Analyses

A Dionex Ultimate 3000 RSLCnano UHPLC system coupled to a Q-Exactive benchtop Orbitrap Mass Spectrometer with Easy Spray Ion Source was used for analyses (Thermo Fisher Scientific, Hemel Hempstead, UK). For all samples, peptides (1 μl corresponding to 1 μg tryptic peptides) were loaded onto a 300 μm ID x 5 mm long 100 Å, 5 µm C_18_ PepMap trap column (Thermo Fisher Scientific, Hemel Hempstead, U) at a flow rate of 8 μl/ min for 5 min in 0.1% (v/v) formic acid in water, then separated using a PepMap RSLC C18 Easy Spray Column (2um, 100A, 75um x 500mm, Thermo Scientific) and binary buffer system of 0.1% (v/v) Formic Acid in Water (Buffer A) and 80% (v/v) Acetonitrile with 0.1% Formic Acid (v/v) at a flow rate of 250nl/min (All Optima LC-MS Grade, Fisher Scientific, Loughborough, UK). Gradients for both Data Dependent Analyses (DDA) and Data Independent Analyses (DIA) were as follows: 2% Buffer B (0-4mins), 2-40% Buffer B (4-100mins), 40-98% Buffer B (100-105mins), 98% Buffer B (105-115mins), 98-2% Buffer B (115-117mins) followed by column equilibration at 2% Buffer B before the next injection.

For DDA analyses, peptides were analyzed with one full scan (400 – 1600 m/z) at 70,000 mass resolution (200m/z), AGC target of 3 x 10^6^ ions and injection time (IT) of 128ms, followed by 15 data-dependent HCD scans (Resolution: 17,500 at 200 m/z, AGC: 1 x 10^5^ ions, max IT: 25ms, Normalized Collision Energy (NCE): 27%, isolation width: 1.5m/z, intensity threshold: 1.2 x 10^5^) with dynamic exclusion enabled (20s). For DIA analyses settings were taken from Sinitcyn et al.^53^ with peptides analyzed using one full scan (350-1400 m/z at 70,000 resolution, AGC target 3 x 10^6^ ions) followed by 48 data-independent MS/MS scans within the mass range of 350-975 m/z at 17,500 resolution (AGC: 3 x 10^6^ ions, max IT: 22ms, isolation width 14m/z, NCE: 25%).

### Media Water Enrichment Analyses via GC-MS

Media water enrichment was analyzed using GC-MS (Trace 1300-ISQ, Thermo Fisher Scientific, Hemel Hempstead, UK) and the approach described in Wilkinson et al.^54^, which is a modification of the protocol described by Mahsut et al.^55^. Briefly, 50 μl of media samples from each well were incubated with 2 μl of 10 M Sodium Hydroxide and 1 μl of acetone for 24 h at room temperature; this high pH incubation leads to the exchange of deuterium from water with the hydrogen positions on the acetone. Following incubation, the acetone was extracted into 200 μl of n-heptane, and 0.5 μl of the heptane phase was injected into the GC for analysis. A standard curve of known D_2_O enrichment (from 0-20 Atom Percent) was run alongside the media samples for calculation of enrichment.

### Myotube Morphology of Cultured Cells

Light microscope images of myotubes were taken after 24h and 48h, for measurement of myotube width and validation of treatment effects. Mean myotube width was calculated from the measurement of 200 myotubes across 10-15 images for each treatment using ImageJ software (National Institutes of Health, Frederick, MD, USA). Data were analysed (using GraphPad Prism v.10 La Jolla, CA, USA) by 1-way ANOVA, with Tukey’s multiple comparison test used to examine differences between treatment groups (*P*<0.05 was considered as statistically significant).

### Biological Function and Gene Ontologies of Significantly Changing Protein FSR

In order to gain biological context for the proteins which fractional synthesis rates were significantly changing, due to either treatment with IGF-1 or DEX, functional enrichment and network analyses were performed. Accession numbers associated with significantly changing proteins were mapped to their gene IDs and functional enrichment of key gene ontologies associated with the significantly changing proteins were determined using the clusterProfiler package^30^, providing information on the top biological functions that may be being affected due to the treatments. Data were visualized as dotplots, gene-concept networks and enrichment maps using the enrichplot^56^ R package.

### Benchmark, software versions

d2ome-DIA was compared to d2ome (v1.5.10). DIA-NN (V1.9.2), Mascot database search engine (version 2.5), clusterProfileR package^30^, and uniport mouse protein sequence database (December 2024 release) were used to identify peptides from tandem mass spectra data.

## Data Availability

The results of the analyses from DIA and DDA approaches reported in this work are available as Supplementary Data. The mass spectral data and data analysis results have been deposited in the MassIVE repository (http://massive.ucsd.edu). The dataset identifier is MSV000099963.

## Code availability

The software for protein turnover determination from DIA approach is available on GitHub https://github.com/rgsadygov/d2ome-DIA.

