## Supplementary Information for "Proteostasis Modelling using Deuterated Water Metabolic Labeling and Data-Independent Acquisition Mass Spectrometry"

### TABLE OF CONTENTS

**Supplementary Note** …………………………………………………………………..….4-7

**Supplementary Figures**

1. Images of C2C12 differentiated myotubes…………………………………………........8
2. The Venn diagrams of common and unique proteins in DDA and DIA methods …………………………………………………………………………………9
3. The comparisons of DDA and DIA quantified peptides in control experiments….10
4. The comparisons of DDA and DIA quantified peptides in dexamethasone (DEX) samples………………………………………………………………………………11
5. The comparisons of DDA and DIA quantified peptides in insulin-like growth factor 1 (IGF-1) treated samples …….……………………………………………………13
6. The pie charts of the percentages of peptides quantified from dexamethasone (DEX) and insulin-like growth factor 1 (IGF-1) treated samples at various levels of goodness-of-fit in DDA and modes using MS1 and MS/MS quantifications…………………….………… …………………………………….……15
7. The comparison of the peptides' turnover rate determined from data-dependent acquisition (DDA) and MS1 from data-independent acquisition (DIA) ……………16

8. The relative difference of the peptides' turnover rate determined from data-dependent acquisition (DDA) and MS1 from data-independent acquisition (DIA) ……………………………………………………………………………….17
9. The comparison of peptide turnover rates from data-dependent acquisition (DDA) and from MS/MS of data-independent acquisition (DIA). …………………………18
10. The relative difference of peptide turnover rates from data-dependent acquisition (DDA) and from MS/MS of data-independent acquisition (DIA)……………………19
11. Volcano plots of turnover rate fold changes between control and dexamethasone (DEX) and IGF-1-treated samples …………………………………………………...20
12. Functional enrichment analyses of insulin-like growth factor 1 (IGF-1) treated samples in data-dependent acquisition (DDA) experiments……………………..…22
13. Functional enrichment analyses of dexamethasone (DEX) treated samples in data-dependent acquisition (DDA) experiments…………………………………….24
14. Functional enrichment analysis of dexamethasone (DEX) treated cells (up regulated in fractional synthesis rate proteins, FSR) in the data dependent acquisition (DDA) mode. ……..……………………………………………………….26
15. Functional enrichment analysis of dexamethasone (DEX) treated cells (proteins downregulated in fractional synthesis rate, FSR) in the data dependent acquisition (DDA) mode.…………………………………………………………………….28
16. Relative turnover rate differences of C2C12 cell proteins in DDA and DIA methods Cambridge and colleagues' study1 using SILAC2 for protein labeling……………………………………………………………………………12

17. The proportions of the numbers of mass isotopomers of fragmentation ions observed in tandem mass spectra acquired in data-independent acquisition (DIA) mode. ……………………………………………………………………………...30
18. The excess deuterium enrichment of the peptide “EGFFELIPQDLIK” computed from its ions (the precursor and fragment ions) and averaged deuteration levels……………………………….…………………………………………31
19. The time course of label incorporation values, which are computed from precursor ions (MS1, represented by blue circles) and convolved fragment ions (MS2, represented by red triangles) over different labeling durations…………………………………………………………………………32
20. The graphical user interface (GUI) to input data and parameters for the protein turnover rate estimations……………..…………………………………………33
21. The graphical user interface provides a detailed visualization of the results from protein turnover studies using metabolic D2O-labeling and LC-MS experiments………………………………………………………………………….34
22. The visualization of the chromatographic elution profiles of quantified peptides at each labeling duration from every replicate mass spectrometry run by d2ome-DIA………………………………….…………. ……………………………35
23. Validation figures generated for the peptide LDEAEQLALK$^{2+}$ (MYH8_MOUSE) before the start of the labeling, derived from the “CellCultureDIA_Dynamics_0h1” experiment file………………………………….…………. …………………..36

**Supplementary Table**

1. Database search engine report file format for D2ome-DIA input …………………… 37

### Supplementary Note

#### D2ome-DIA:

D2ome-DIA is a stand-alone software tool designed to compute turnover rates for proteins and peptides from heavy water-labeled, data-independent LC-MS experiments. The software quantifies the abundance of mass isotopomers of peptides and proteins, computes their corresponding turnover rates, and provides a graphical user interface (GUI) for visualizing the quantified relative abundances of these mass isotopomers along with their turnover rates. Additionally, D2ome-DIA includes a validation feature that allows for the analysis of each individual MS1 and MS2 scans used in quantifying the abundance and turnover rates of proteins.

#### Inputs and outputs of the computation tool

The primary inputs for the software are LC-MS spectral files in the "mzML" format and database search results in comma-separated (.csv) file format. D2ome-DIA is compatible with DIA-NN database search outputs and also accepts results from any search engine, provided they are preprocessed as specified in **Supplementary Table 1**. The database search results must contain the following information: peptide sequence, start elution time, end elution time, precursor mass, and a list of the top four fragment ions along with their corresponding quality scores.

The software generates two primary outputs: mass isotopomers abundance quantifications and turnover rate estimations. The quantification results consist of intensity values for precursor and fragment ions, including the mono-isotope and the first five heavy mass isotopomers. These results are provided as comma-separated values

(CSV) in two formats: "Protein.Quant.csv" and "MS2_QuantificationResult.csv." The first file, "Protein.Quant.csv," contains detailed information about each peptide of a protein, along with the intensities of its mass isotopomers quantified from $MS_1$ spectra. The second file, "MS2_QuantificationResult.csv," offers similar information for fragment ions that are quantified from $MS_2$ spectra.

The turnover rate estimation outputs include the degradation rates for proteins and peptides, 95% confidence interval and goodness-of-fit (GOF) measures that compare the experimental data to the theoretical curve fitting. The rates are reported in two forms: “Protein.RateConst.csv,” which contains the turnover rates determined from both $MS_1$ and $MS_2$ spectra for each constituent peptide of a protein, and “Analyzed_Proteins_and_Their_Rates.csv,” which provides a summary of the turnover rates for all proteins identified in the given LC-MS experiment.

In addition, the software generates isotope profiles for each peptide extracted from all MS1 scans used in the quantification process. It also creates isotope profiles for the fragment ions of each peptide based on all MS2 scans, which are utilized for quantification. These profiles, along with their integrated values, are reported as parquet files for each experimental dataset. They cover every peptide and fragment ion listed in the DIA-NN database search results involved in the quantification. These files are essential for validating the elution profile, label enrichment, and other parameters calculated during the rate computation process.

**Visualization**

The d2ome-DIA software features a visualization tab that includes two main charts illustrating the time series of monoisotopic Relative Abundance (RA) values used for calculating the turnover rates of peptides and proteins, as shown in **Supplementary Figure 21**. This tab provides easy access to the turnover rate estimation results for each protein. For every peptide associated with a protein, the monoisotopic RA values are estimated from the isotope profiles and compared to the theoretical fit. This method effectively visualizes the relationship between the experimental data points and the expected theoretical values, which are computed based on the estimated turnover rate.

**Tools for Manual Validation**

d2ome-DIA allows users to inspect the quality of the quantified isotope profiles of the mass isotopomers from spectra files through a comprehensive graphical presentation. The software provides a graphical representation of the peptides' raw and relative abundances of mass isotopomers over the elution window, the amount of label incorporation throughout time course experiments, and detailed spectrum plots for every MS1 and MS2 scan utilized in the quantifications, **Supplementary Figure 21**.

The software utilizes all the intermediate Parquet files generated during the quantification stage to generate the graphical representation presented in the GUI. These files contain detailed information about every MS1 and MS2 scan used in the quantification process. Specifically, they include information about the precursor and fragment ions, such as sequence, charge state, and mass-to-charge ratio. Additionally, the files contain the scan number, retention time, raw abundance of the monoisotope and the first five heavy mass isotopomers, and the amount of label incorporation calculated from the isotope profile.

**Supplementary Figures 22 and 23** present the validation data generated for the peptide LDEAEQLALK$^{2+}$ (MYH8_MOUSE) at zero days of labeling, derived from the “CellCultureDIA_Dynamics_0h1” experiment file. Figures 4A and 4B depict a scatter plot of the raw abundance of the monoisotope and the percentage of label incorporation over the elution window of the peptide.

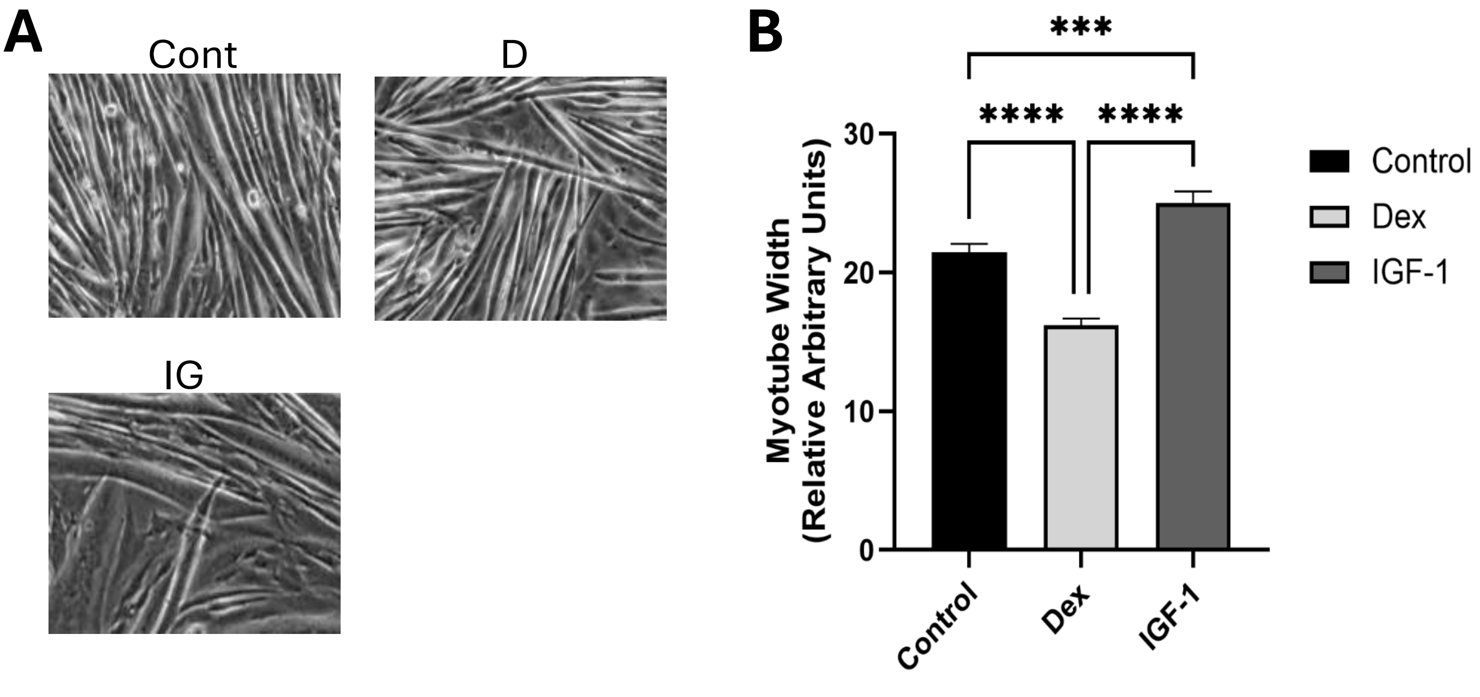


**Supplementary Figure 1.** Images of C2C12 differentiated myotubes (**A**) and myotube width measurements (**B**) after 24h dexamethasone (DEX) or insulin-like growth factor 1 (IGF-1) treatment. Light microscope images were taken from differentiated myotubes following 24h of 50 ng/ml IGF-1 treatment or 2.5 μM Dex treatment. Images were used to calculate mean myotube diameter. Data are expressed as mean + SEM (standard error of the mean, relative arbitrary units).

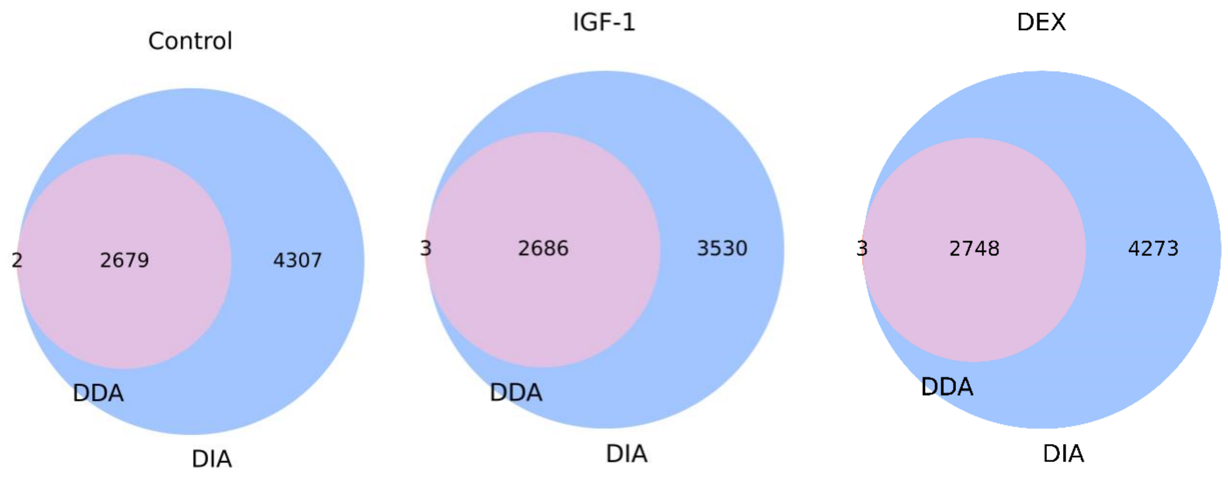


(**A**) (**B**) (**C**)

**Supporting Figure 2.** The number of common and unique proteins quantified in d2ome-DDA and d2ome-DIA in **A**) Control, **B**) insulin-like growth factor 1 (IGF-1)-treated samples, **C**) dexamethasone (DEX)-treated samples. DDA – data-dependent acquisition, DIA-data-dependent acquisition.

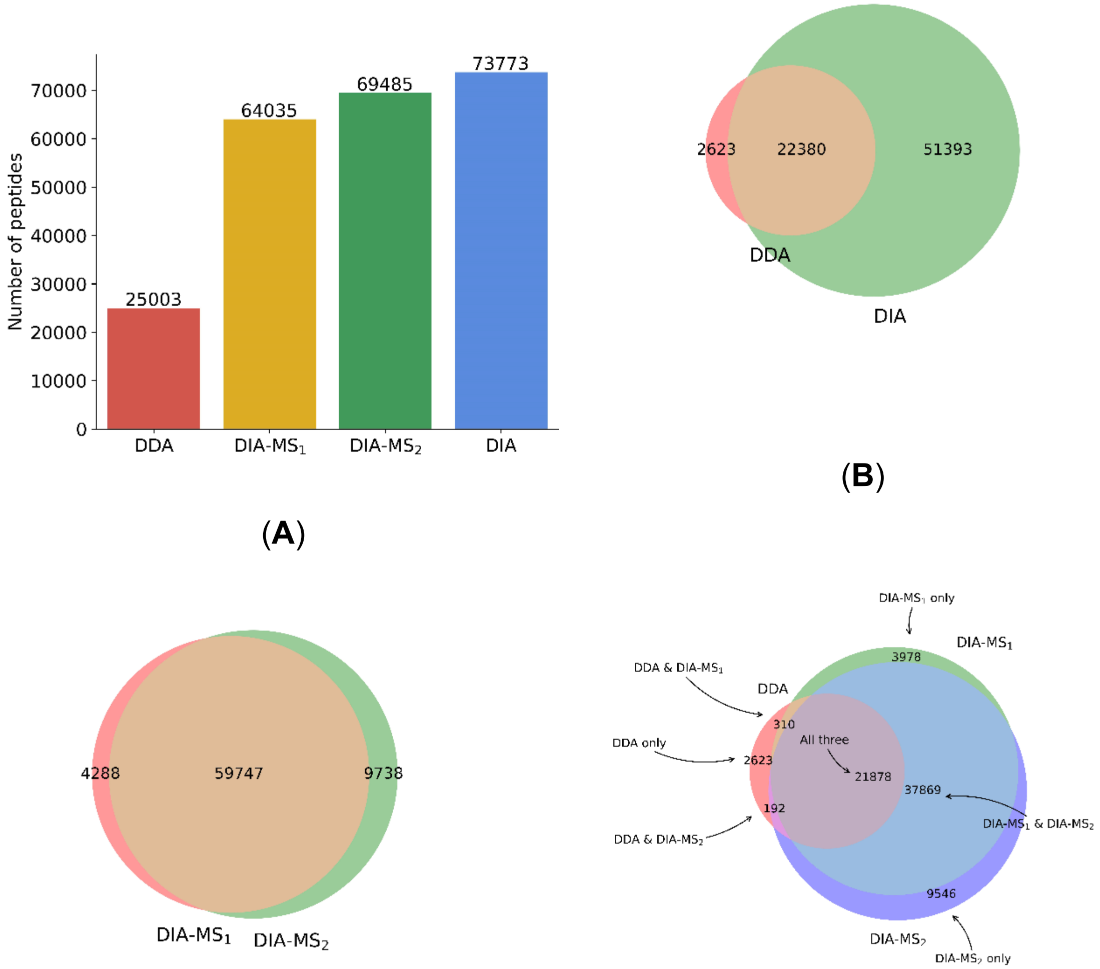


(**A**) (**B**) (**C**) (**D**)

**Supplementary Figure 3.** The number of peptides quantified from the control experiment **A**) Comparison of the number of peptides quantified using d2ome-DDA, d2ome-DIA from MS1 spectral only (DIA-MS1), d2ome-DIA from tandem mass spectra (MS2) spectral only (DIA-MS2), and d2ome-DIA from both MS1 and MS2 spectral **B**) Comparison of the number of peptides quantified using d2ome-DIA and d2ome-DDA. Comparison of the number of peptides quantified using d2ome- DDA and d2ome- DIA MS1 spectra only. **C**) Comparison of the number of peptides quantified using d2ome- DDA and d2ome- DIA

MS2 spectra only. **D**) Three-way Venn diagram showing the overlap of peptides identified by DDA, DIA-MS1, and DIA-MS2 in the control condition.

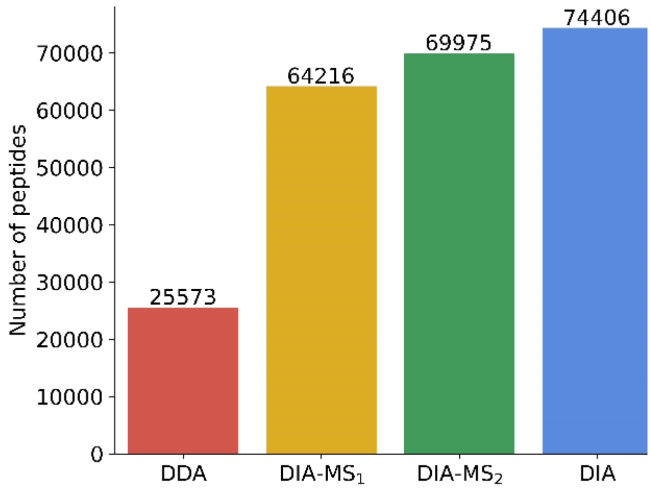


(**A**)

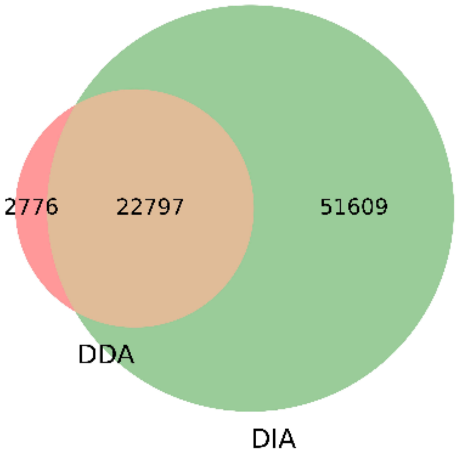


(**B**)

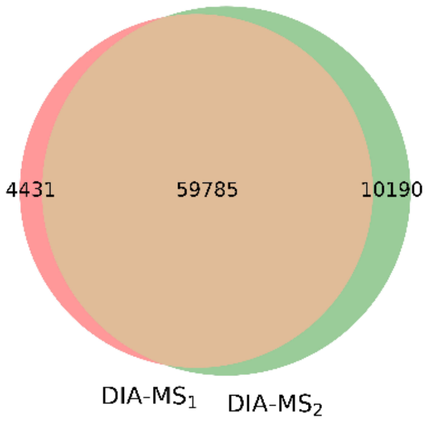


(**C**)

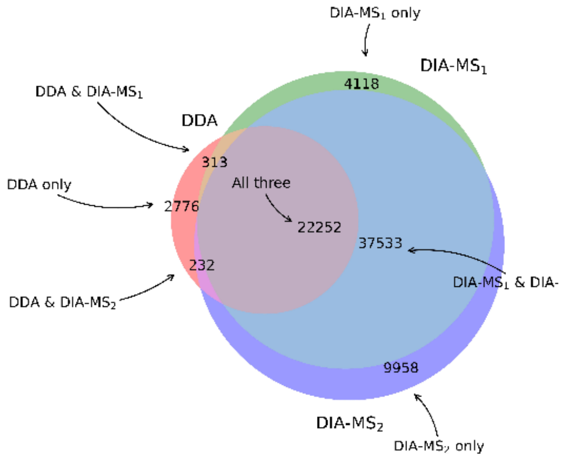


(**D**)

**Supplementary Figure 4.** The number of peptides quantified from the DEX treated samples **A**) Comparison of the number of peptides quantified using d2ome-DDA, d2ome-DIA from only the MS1 spectra (DIA-MS1), d2ome-DIA from only the tandem mass, MS2, spectra (DIA-MS2), and d2ome-DIA from both MS1 and MS2 spectra. **B**) Comparison of the number of peptides quantified using d2ome-DIA and d2ome-DDA. Comparison of the number of peptides quantified using d2ome- DDA and d2ome- DIA MS1 spectra only. **C**) Comparison of the number of peptides quantified using d2ome- DDA and d2ome- DIA MS2 spectra only. **D**) Three-way Venn diagram showing the overlap of peptides identified by DDA, DIA-MS1, and DIA-MS/MS in the DEX experiment.

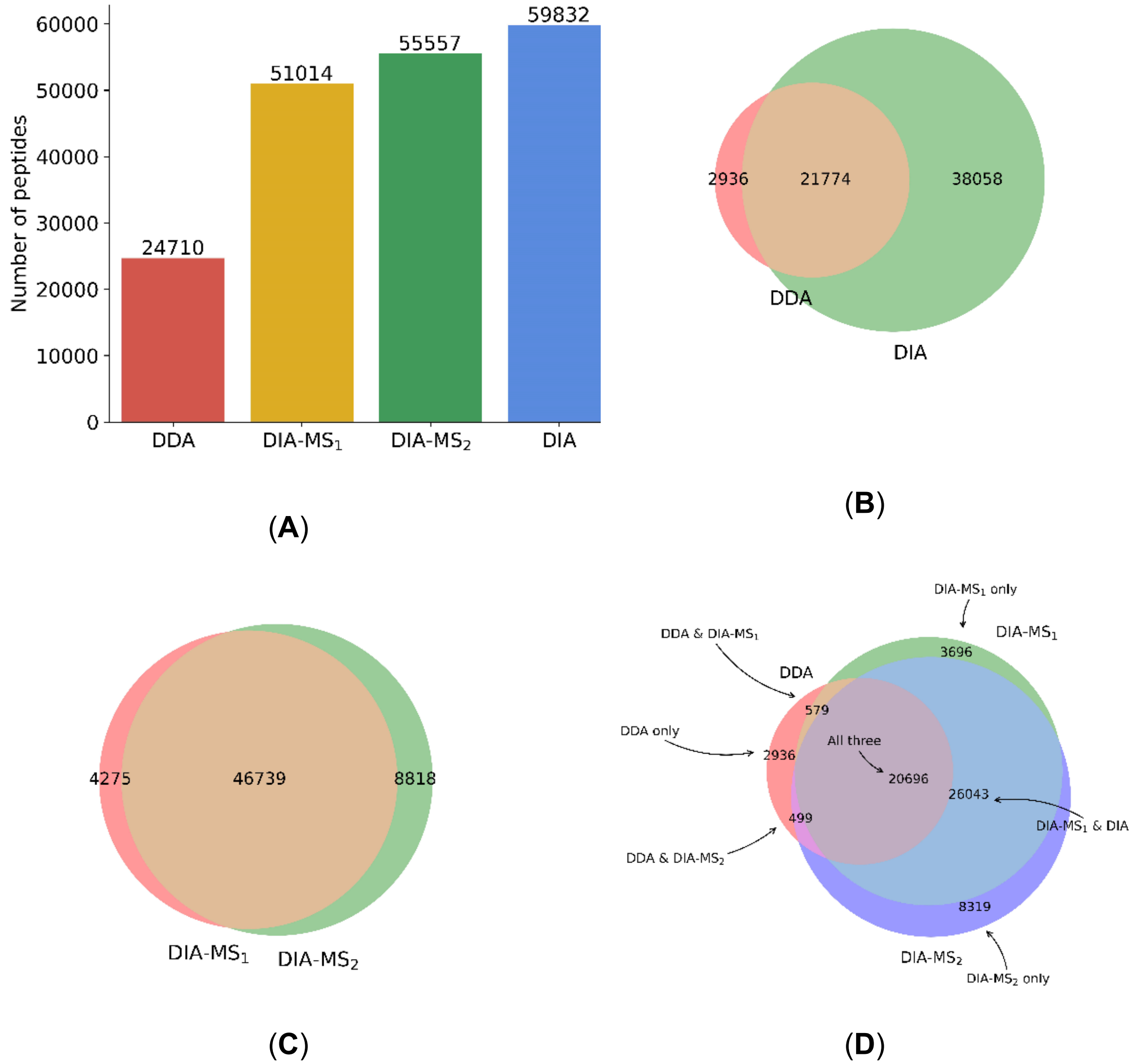


**Supplementary Figure 5.** The number of peptides quantified from the IGF experiment **A**) Comparison of the number of peptides quantified using d2ome-DDA, d2ome-DIA from only the MS1 spectral (DIA-MS1), d2ome-DIA from only the tandem mass, MS2, spectra (DIA-MS2), and d2ome-DIA from both MS1 and MS2 spectra, **B**) Comparison of the number of peptides quantified using d2ome-DIA and d2ome-DDA. Comparison of the number of peptides quantified using d2ome- DDA and d2ome- DIA MS1 spectra only. **C**)

Comparison of the number of peptides quantified using d2ome- DDA and d2ome- DIA MS/MS spectra only. **D**) Three-way Venn diagram showing the overlap of peptides identified by DDA, DIA-MS1, and DIA-MS/MS in the IGF experiment.

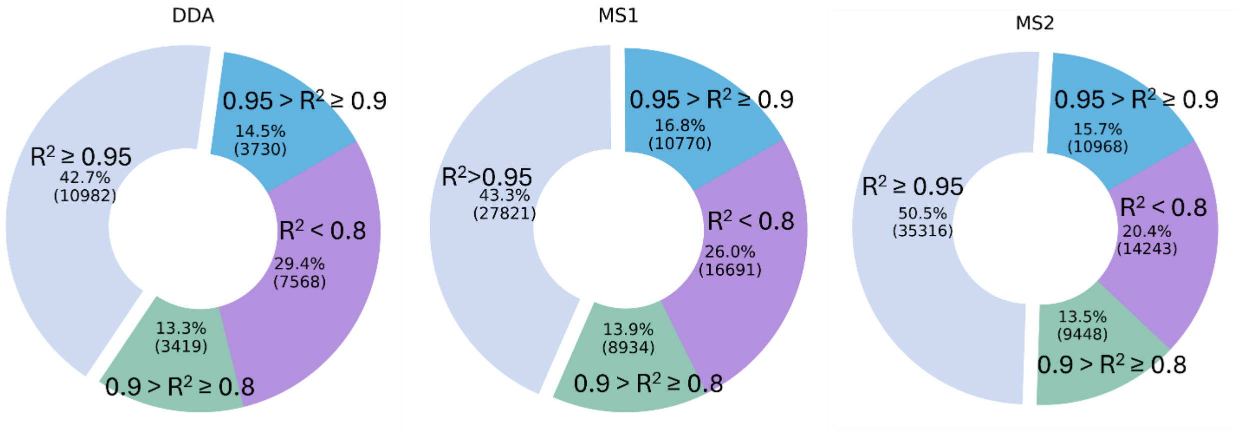


**(A)**

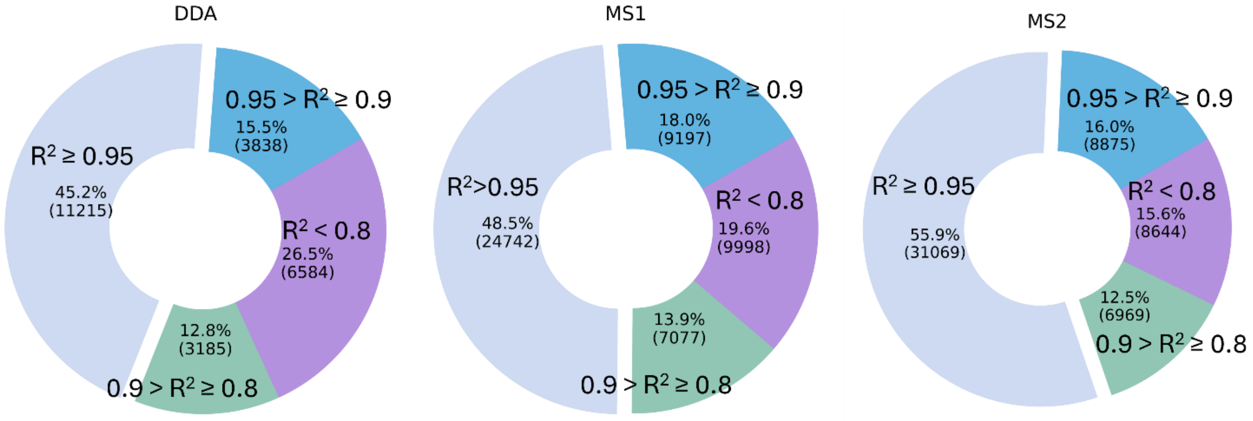


**(B)**

**Supplementary Figure 6**. The pie charts of the percentages of peptides quantified from dexamethasone **A)** (DEX)-treated and insulin like growth factor 1 (IGF-1)-treated **B**) samples at various levels of goodness-of-fit (expressed by the coefficient of determination, $R^2$) in data-dependent (DDA) mode, data-independent (DIA) mode using MS1 quantification, and DIA mode using MS/MS quantification.

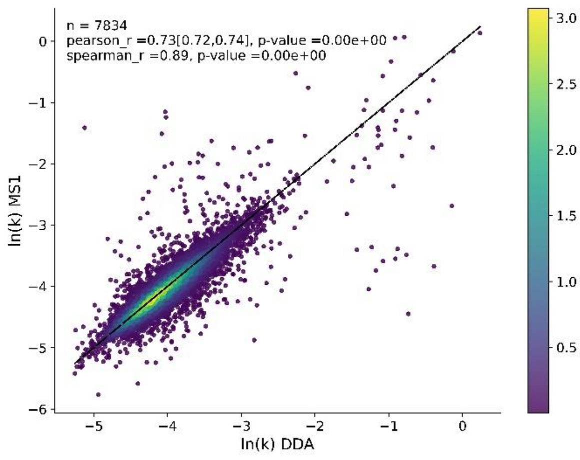


**(A)**

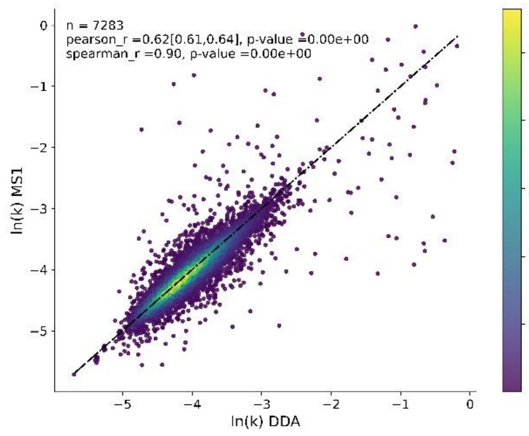


(**B**)

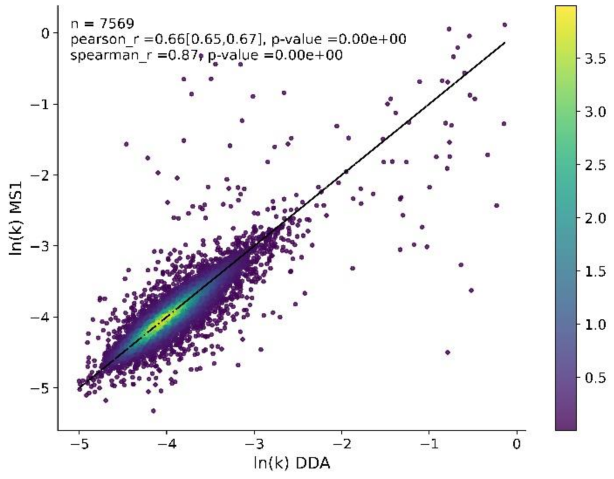


(**C**)

**Supplementary Figure 7**. Scatter plots for the logarithms of peptide turnover rates from data-dependent acquisition (DDA) vs MS1 from data-independent acquisition (DIA) in the **A**) control, **B**) DEX-treated, and **C**) IGF1-treated samples. The peptides were filtered for $R^2 \geq 0.95$ and NDP (number of data points) ≥4.

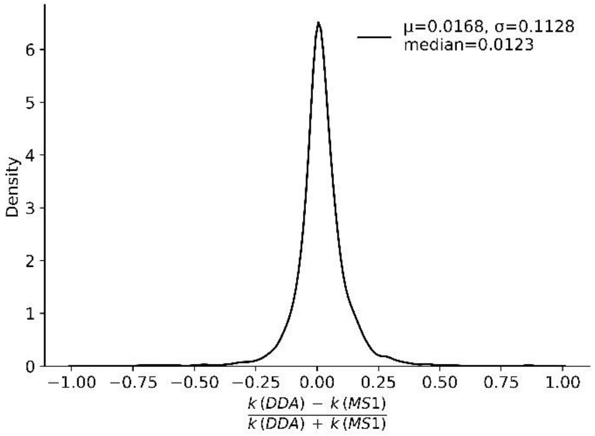


**(A)**

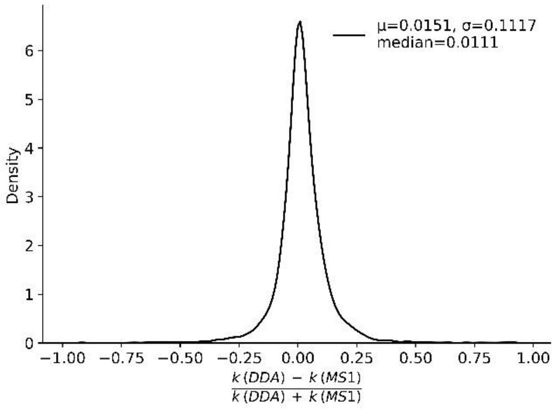


(**B**)

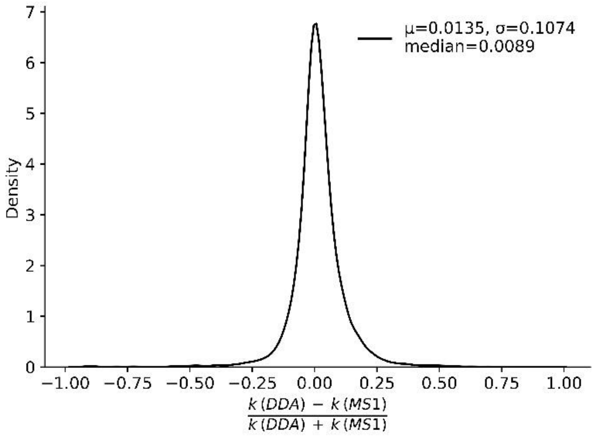


(**C**)

**Supplementary Figure 8**. Density plots of differences of the peptide turnover rates from data-dependent acquisition (DDA) and from MS1 of data-independent acquisition (DIA) in the **A**) control, **B**) DEX-treated, and **C**) IGF1-treated samples. The peptides were filtered for the coefficient of determination ($R^2$) ≥ 0.95 and number of data points (NDP) ≥4.

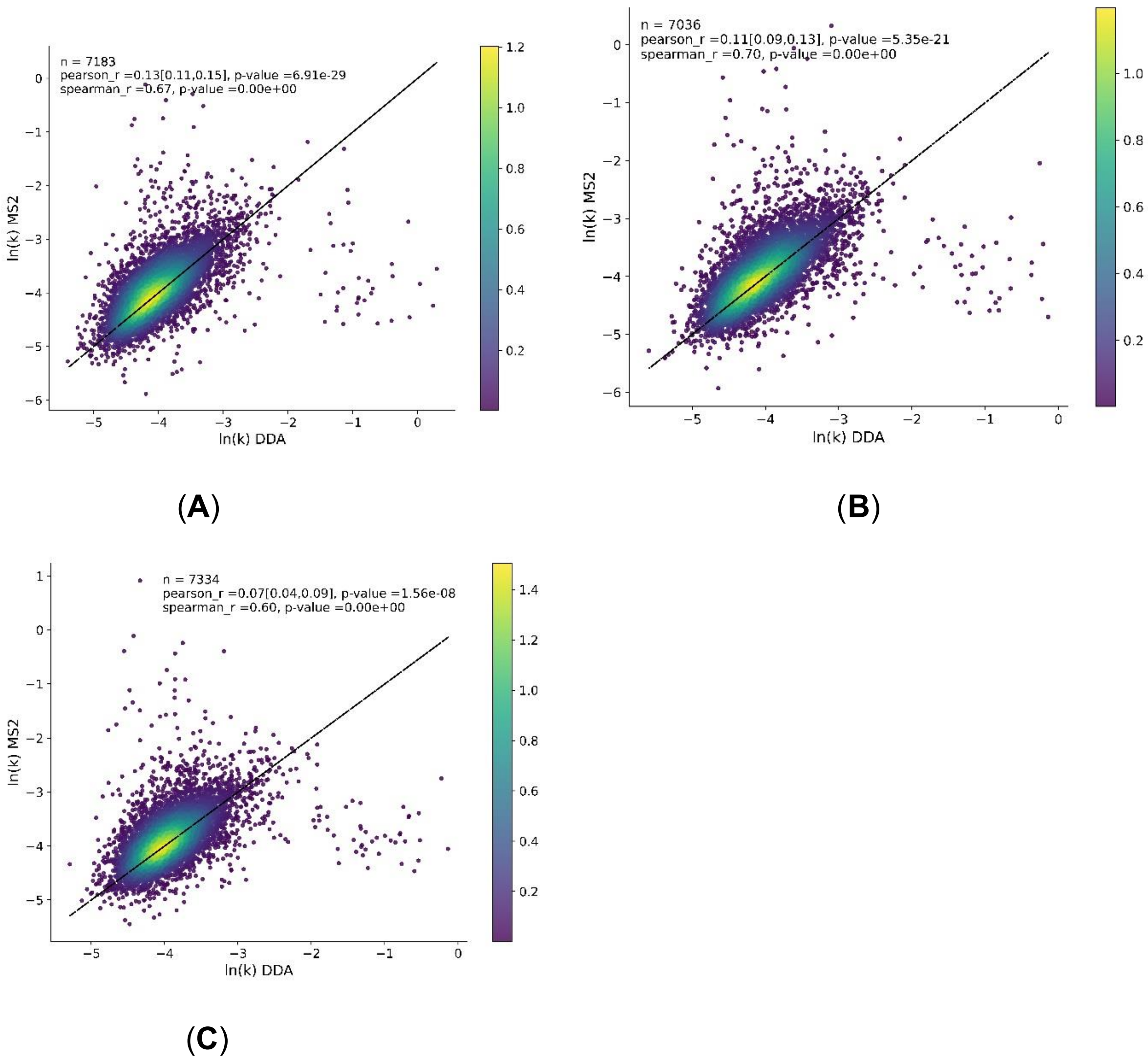


**Supplementary Figure 9**. Scatter plots for the logarithms of peptide turnover rates from data-dependent acquisition (DDA) vs from MS/MS of data-independent acquisition (DIA) in the **A**) control, **B**) DEX-treated, and **C**) IGF1-treated samples. The peptides were filtered for $R^2 \geq 0.95$ and NDP (number of data points) ≥4.

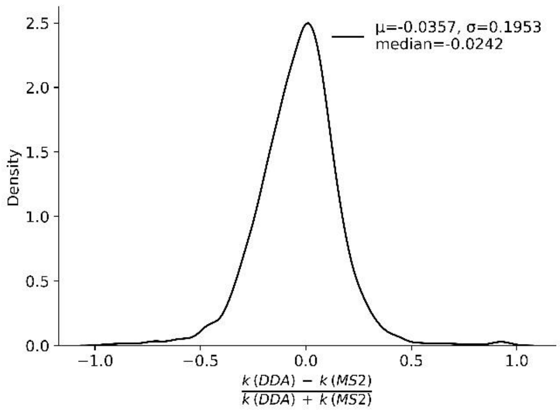


(**A**)

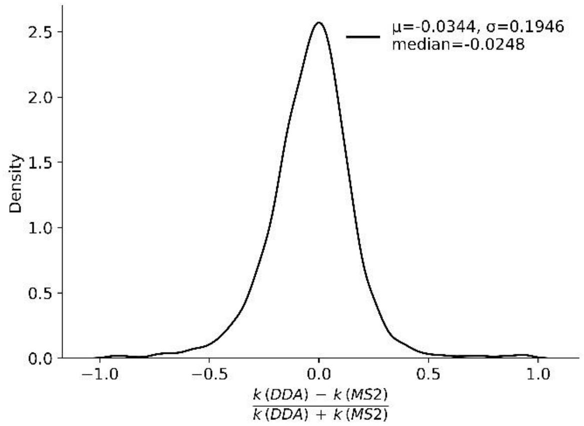


(**B**)

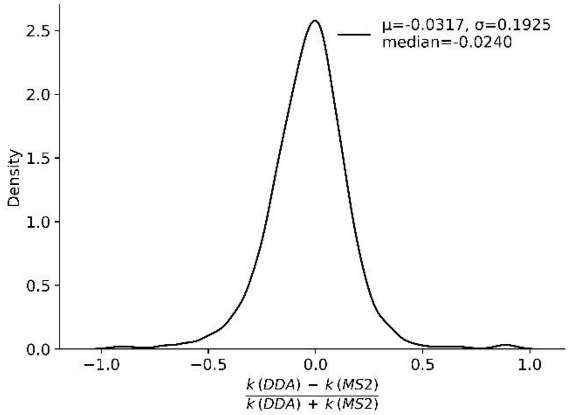


(**C**)

**Supplementary Figure 10**. Density plots of differences of the peptide turnover rates from data-dependent acquisition (DDA) and from MS/MS of data-independent acquisition (DIA) in the **A**) control, **B**) DEX-treated, and **C**) IGF1-treated samples. The peptides were filtered for $R^2 \geq 0.95$ and NDP (number of data points) ≥4.

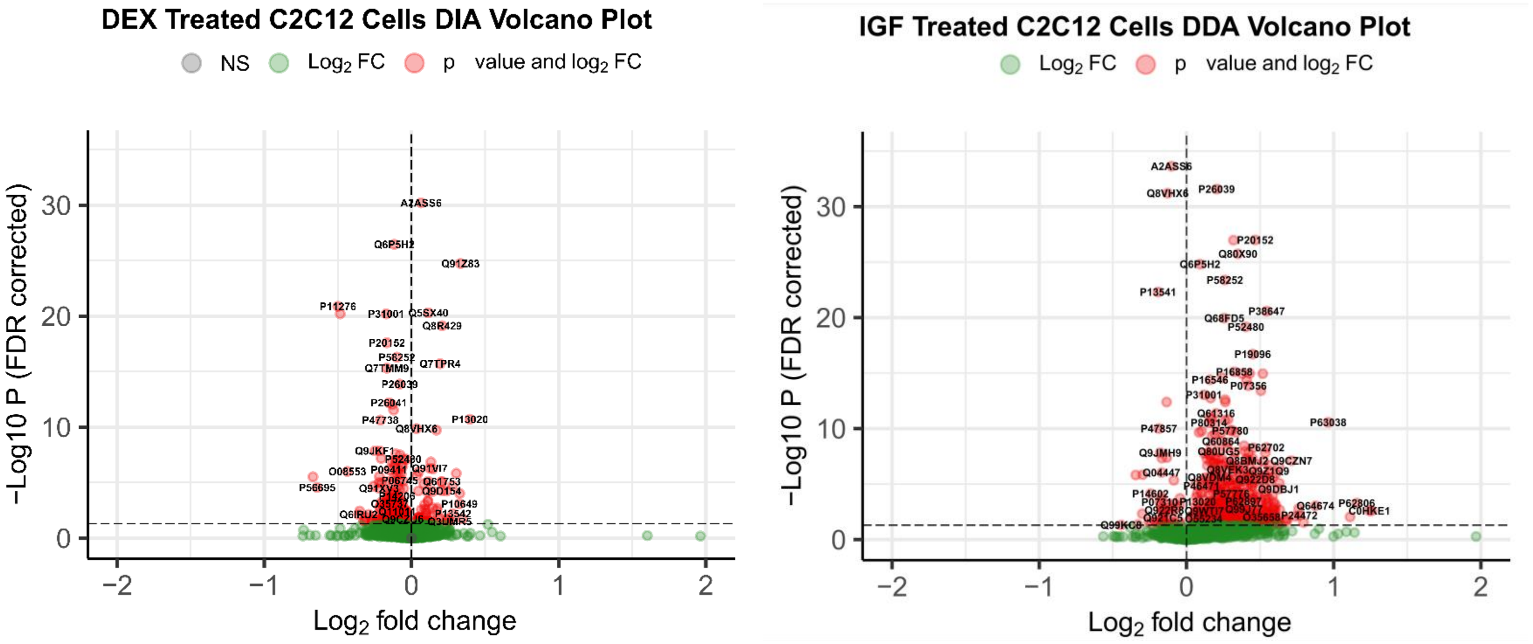


**A** **B**

**Supplementary Figure 11**. Despite the difference in number of proteins detected, the same patterns as in data-independent acquisition (DIA) were present for the data-dependent acquisition (DDA) method. **A**) With dexamethasone (DEX) treatment, there were proteins that both increased and decreased their rates, however the majority of the proteins decreased. **B**) With insulin-like growth factor 1 (IGF-1) treatment, many proteins increased their rates of turnover.

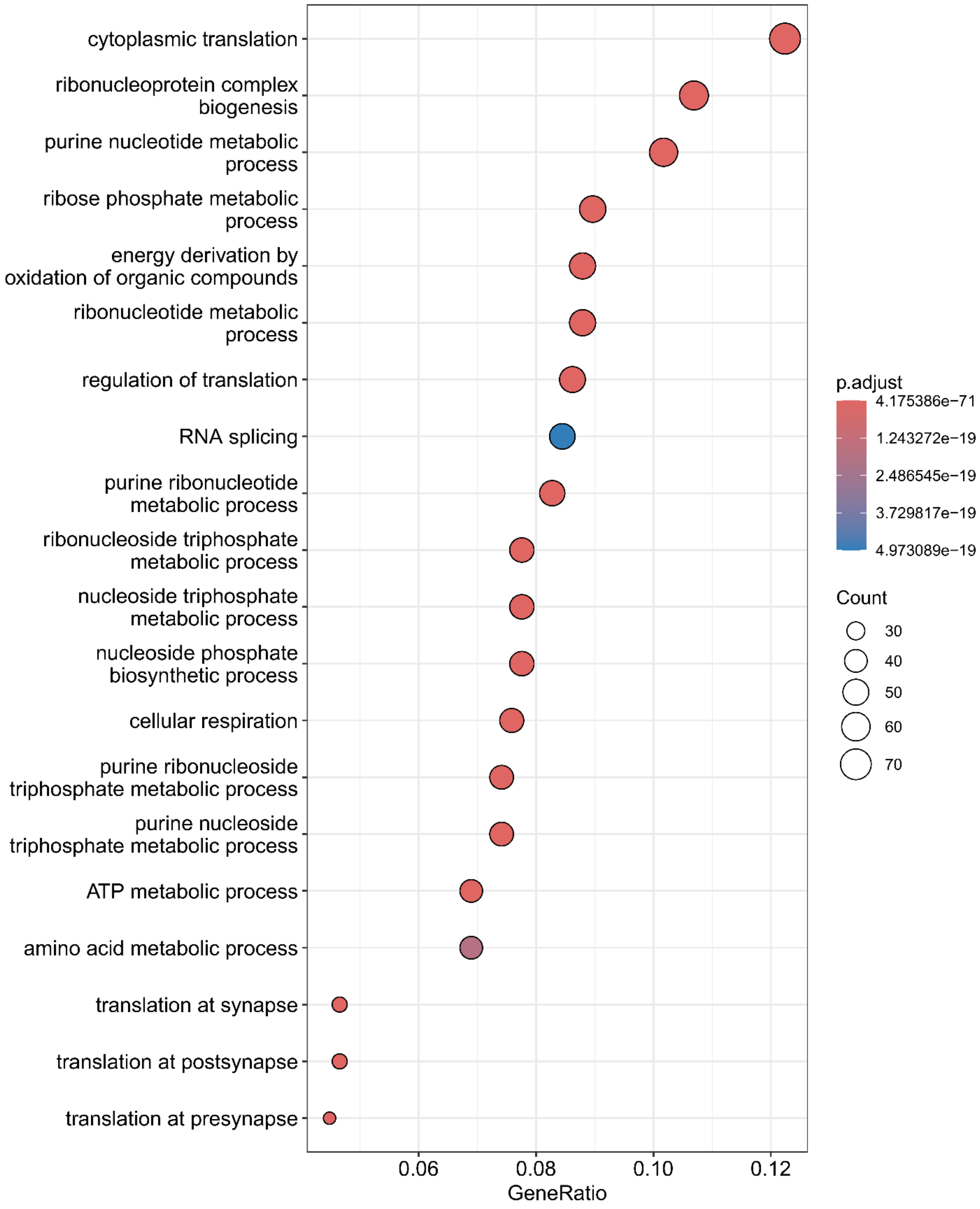


**Supplementary Figure 12 A**

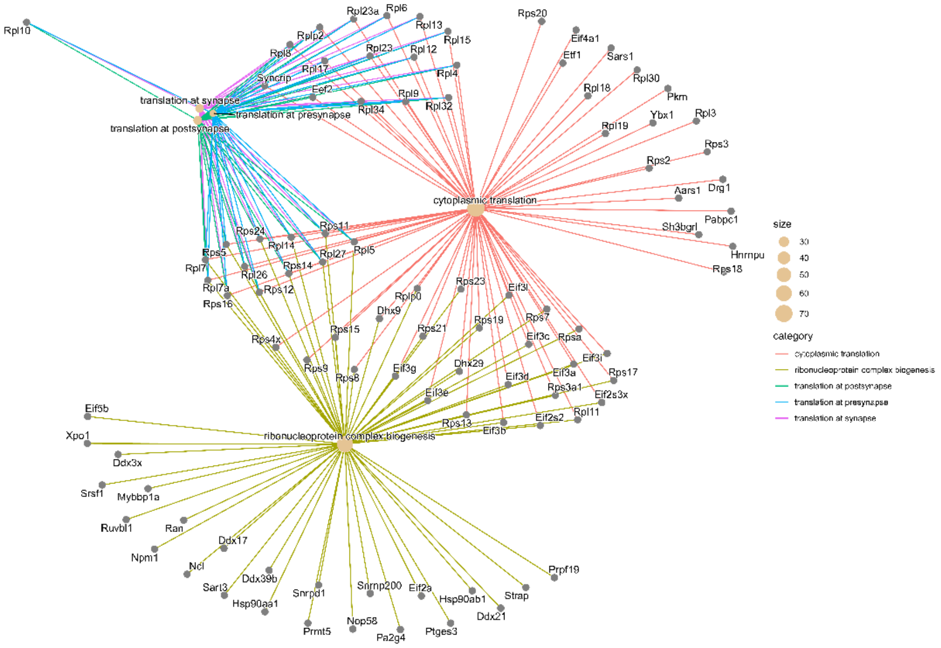


**Supplementary Figure 12 B**

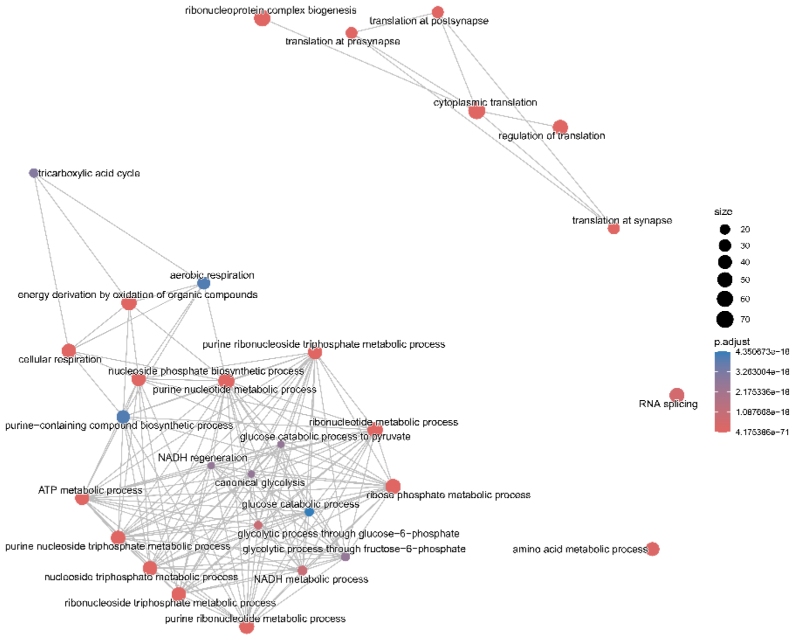


**Supplementary Figure 12 C**

**Supplementary Figure 12.** Functional enrichment analyses of insulin-like growth factor 1 (IGF-1) treated cells in data-dependent acquisition (DDA) mode.

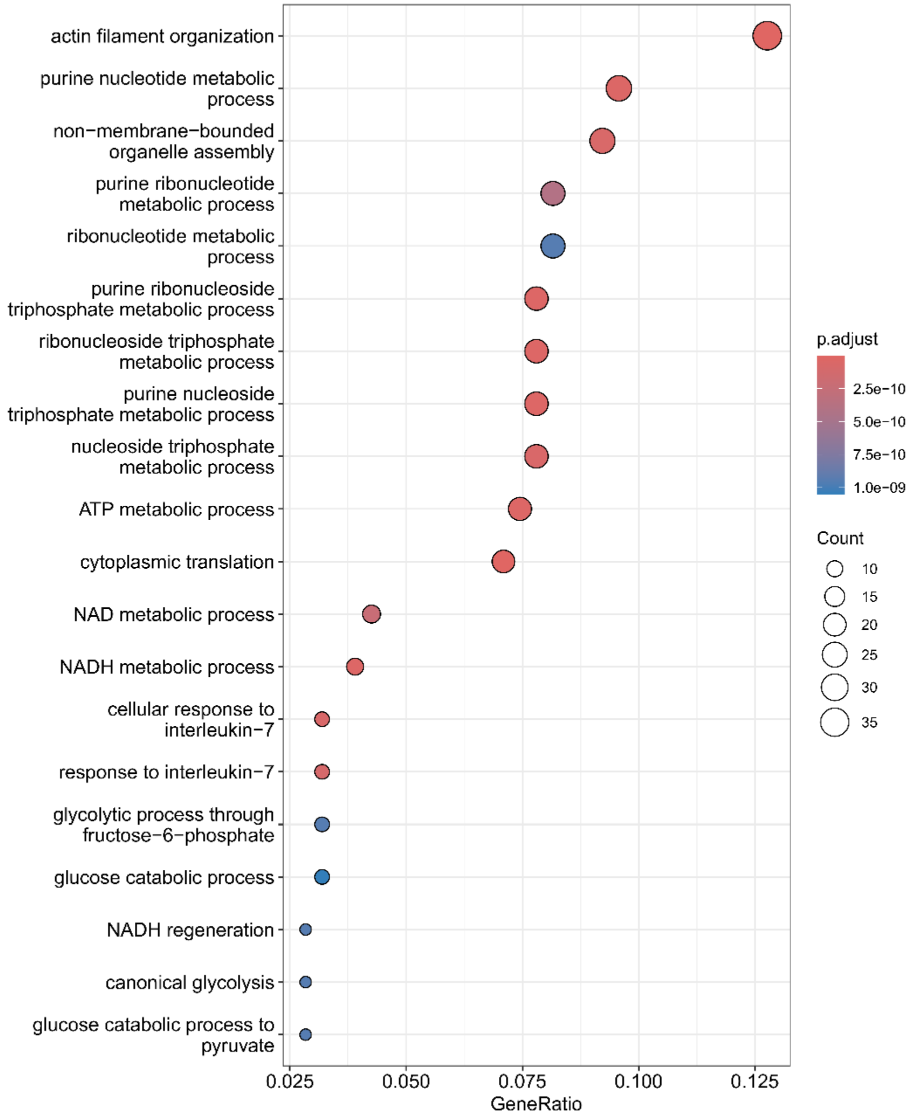


**Supplementary Figure 13 A**

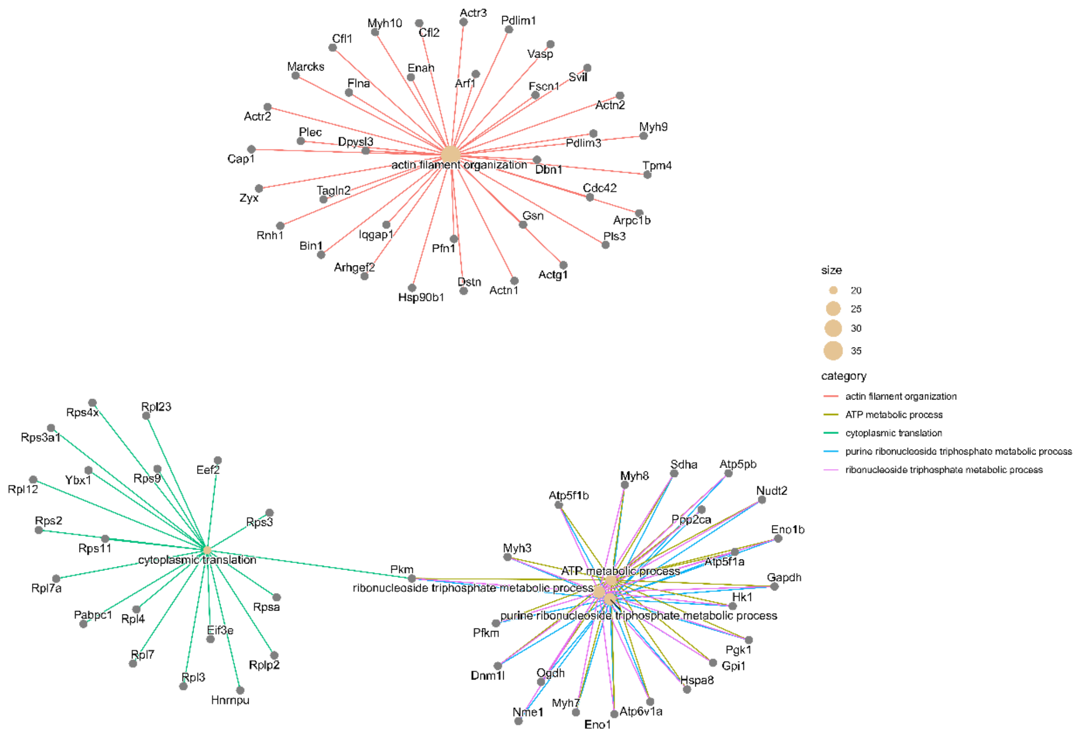


### Supplementary Figure 13 B

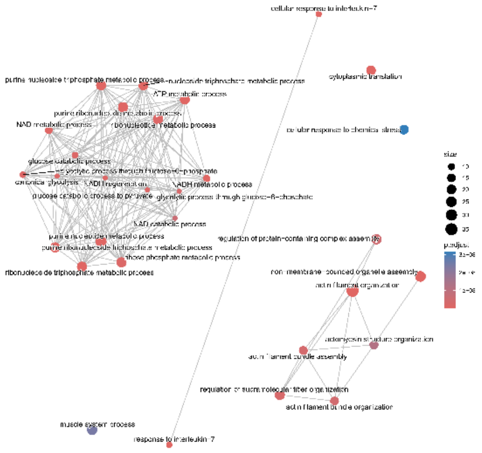

### Supplementary Figure 13 C

**Supplementary Figure 13**. Functional enrichment analyses of dexamethasone (DEX) treated cells (all proteins) in data-dependent acquisition (DDA) mode.

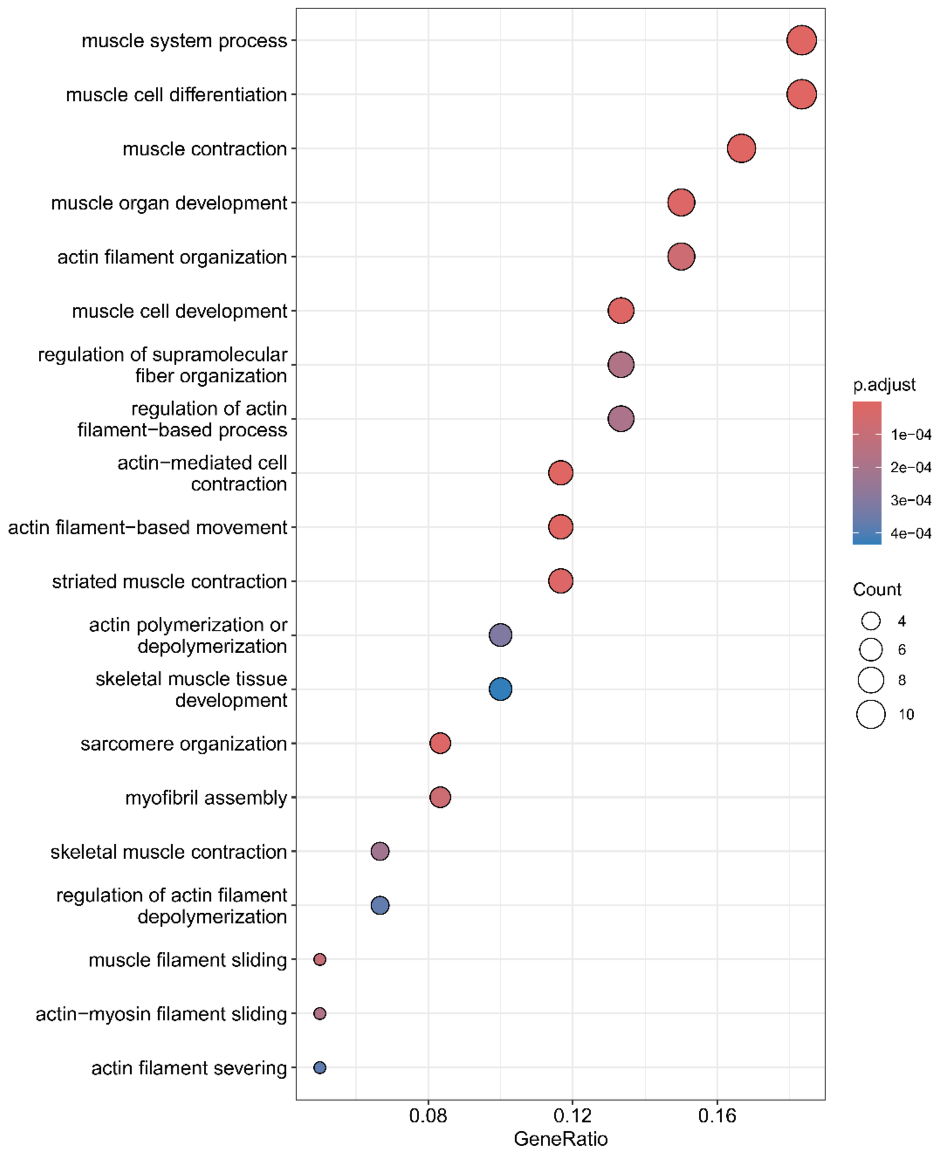


**Supplementary Figure 14 A**

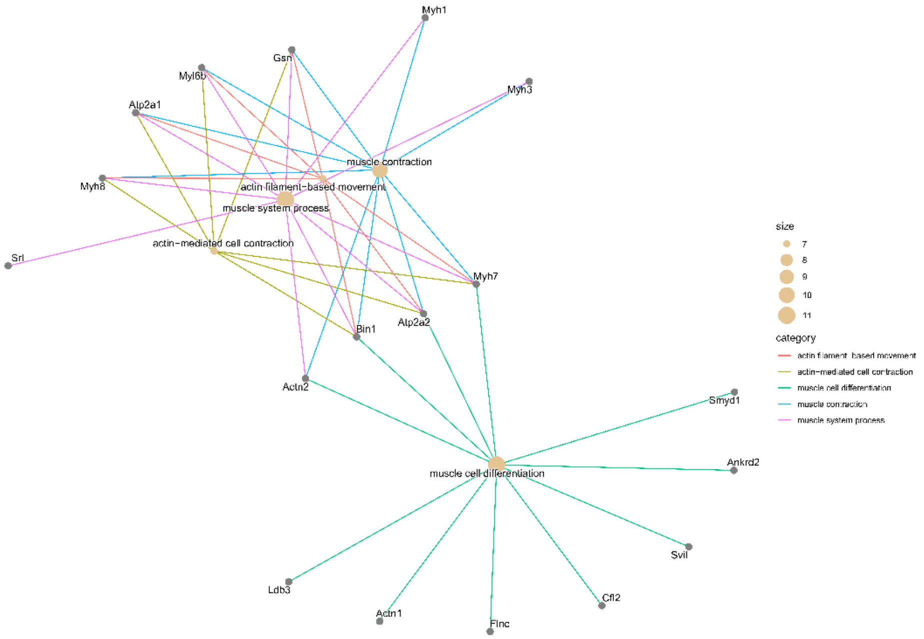


**Supplementary Figure 14 B**

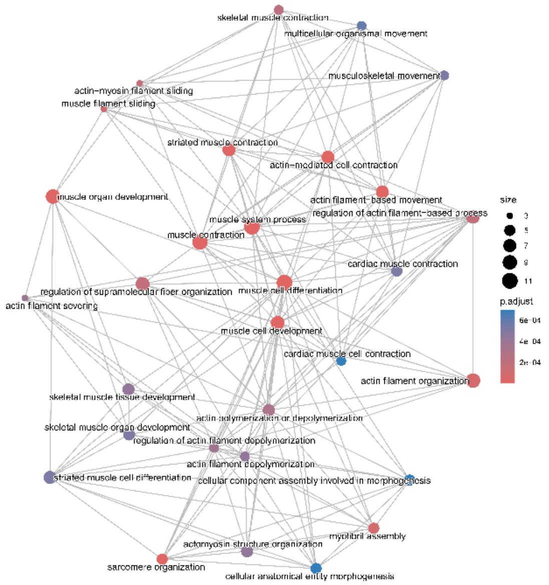


**Supplementary Figure 14 C**

**Supplementary Figure 14**. Functional enrichment analysis of dexamethasone (DEX) treated cells (up-regulated in fractional synthesis rate proteins, FSR) in the data-dependent acquisition (DDA) mode.

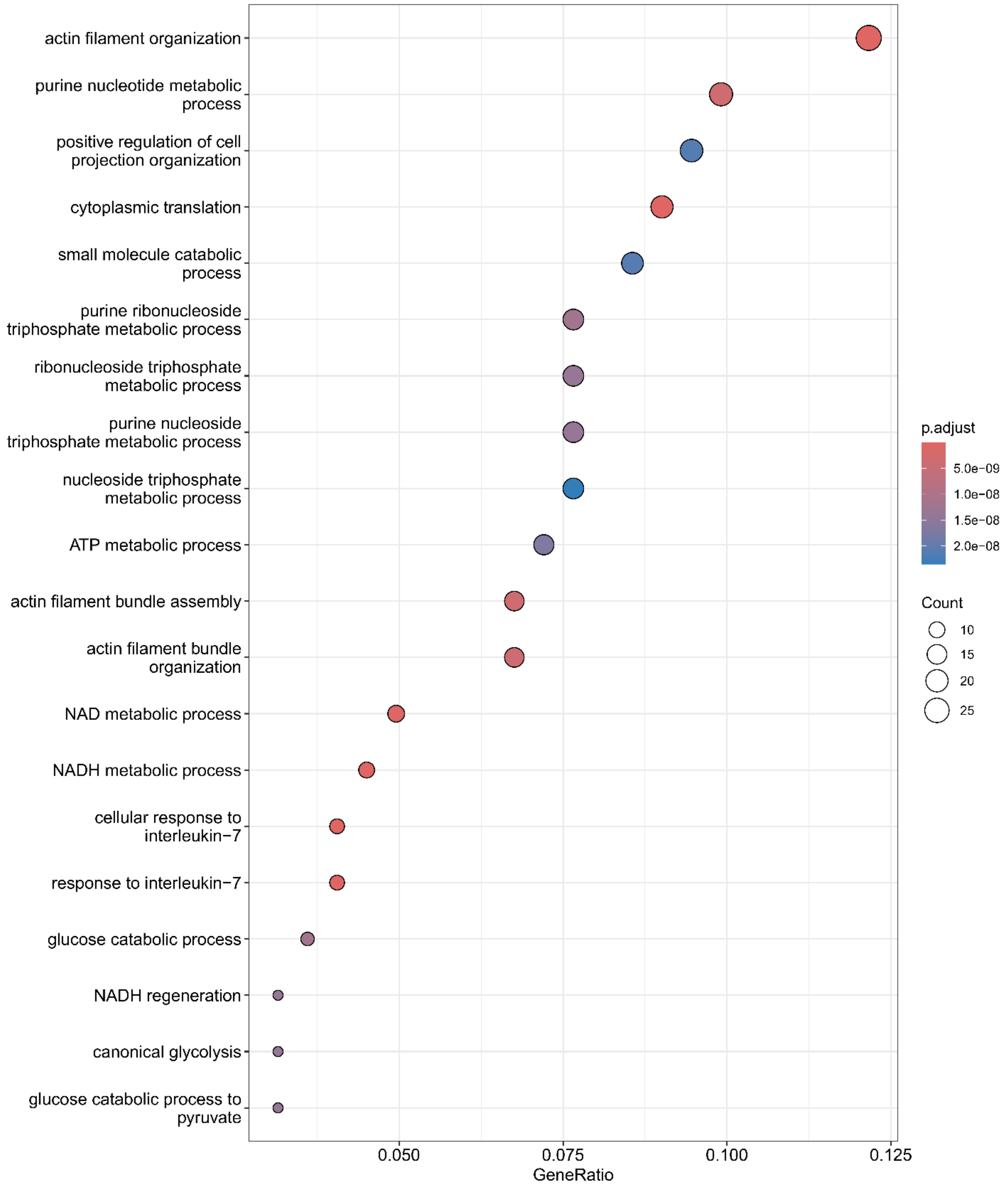


**Supplementary Figure 15 A**

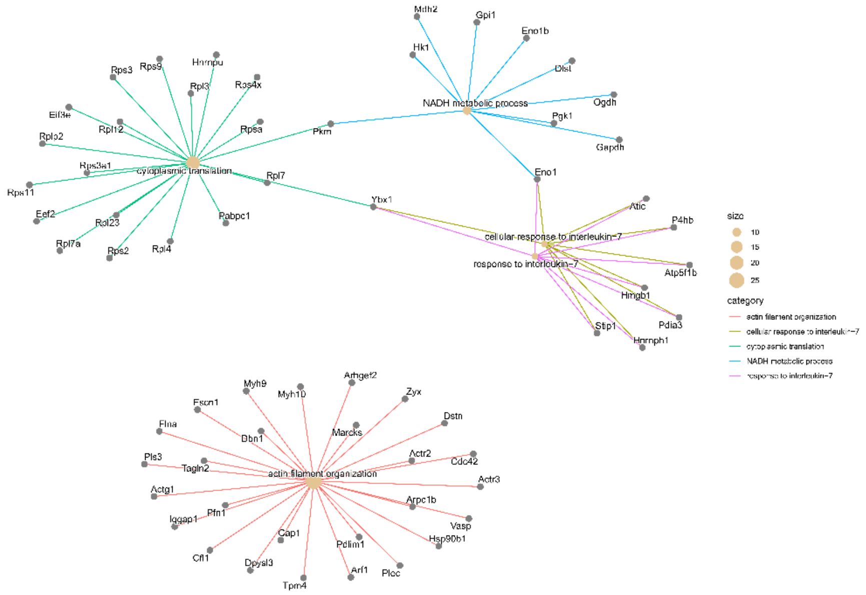


### Supplementary Figure 15 B

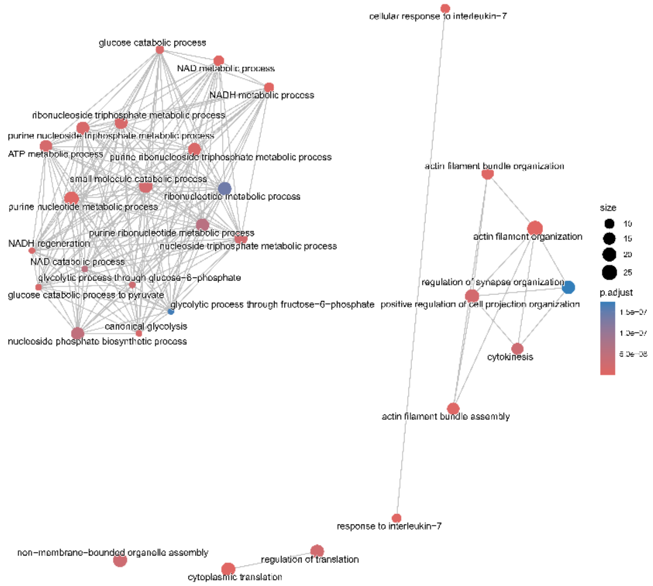


### Supplementary Figure 15 C

**Supplementary Figure 15**. Functional enrichment analysis of dexamethasone (DEX) treated cells (proteins downregulated in fractional synthesis rate, FSR) in the data-dependent acquisition (DDA) mode.

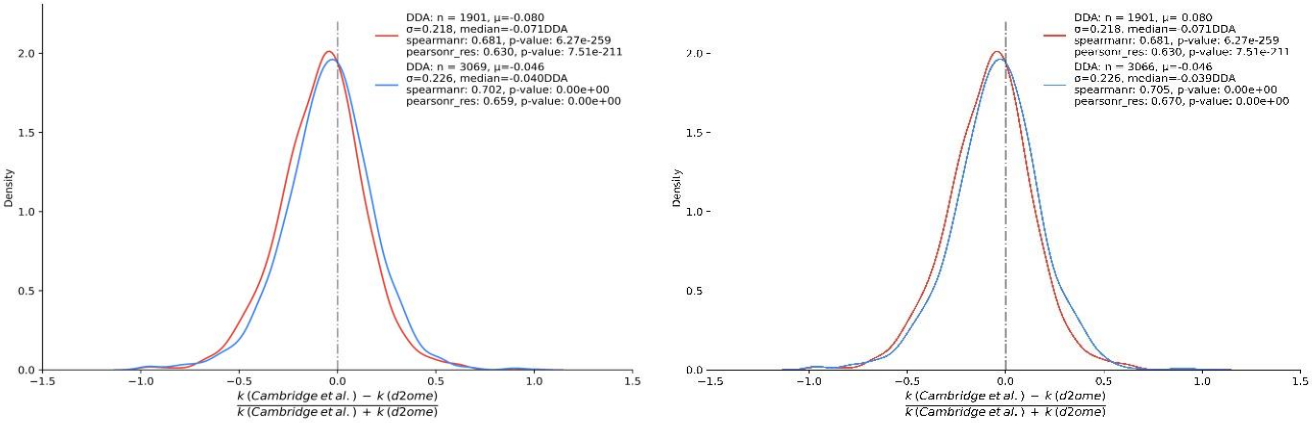


**A)** **B)**

**Supplementary Figure 16**. Density plots of the relative turnover rate differences of C2C12 cell proteins computed in this work with data-dependent (DDA) and data-independent (DIA) methods with those from Cambridge and colleagues' study[1] using SILAC for protein labeling. **A**) results from using the linear regression analysis for estimating label enrichment from tandem mass spectra, **B**) the results from the isotope averaging approach to the estimation of label enrichment.

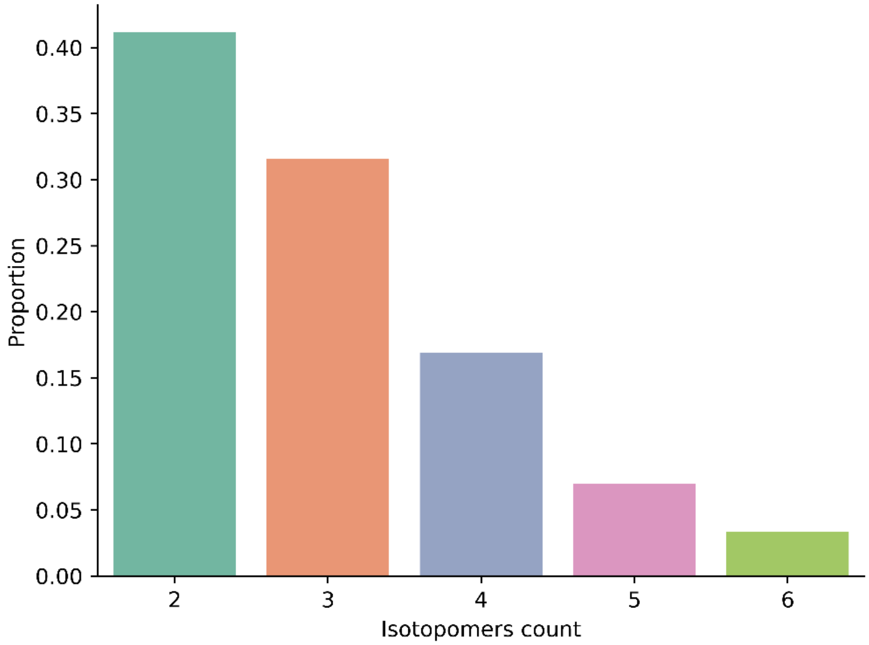


**Supplementary Figure 17**. The bar plot of the proportions of the numbers of mass isotopomers of fragmentation ions observed in tandem mass spectra acquired in data-independent acquisition (DIA) mode.

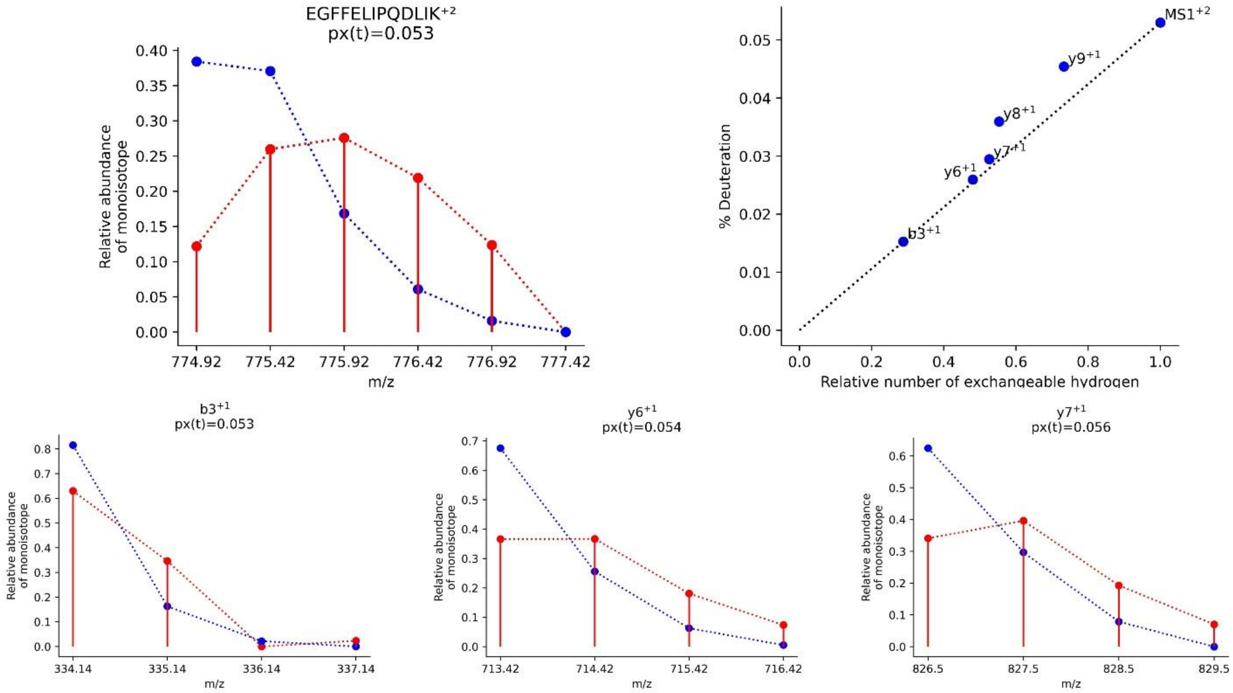


**Supplementary Figure 18**. The excess deuterium enrichment of the peptide "EGFFELIPQDLIK" computed from its ions (the precursor and fragment ions) and averaged deuteration levels.

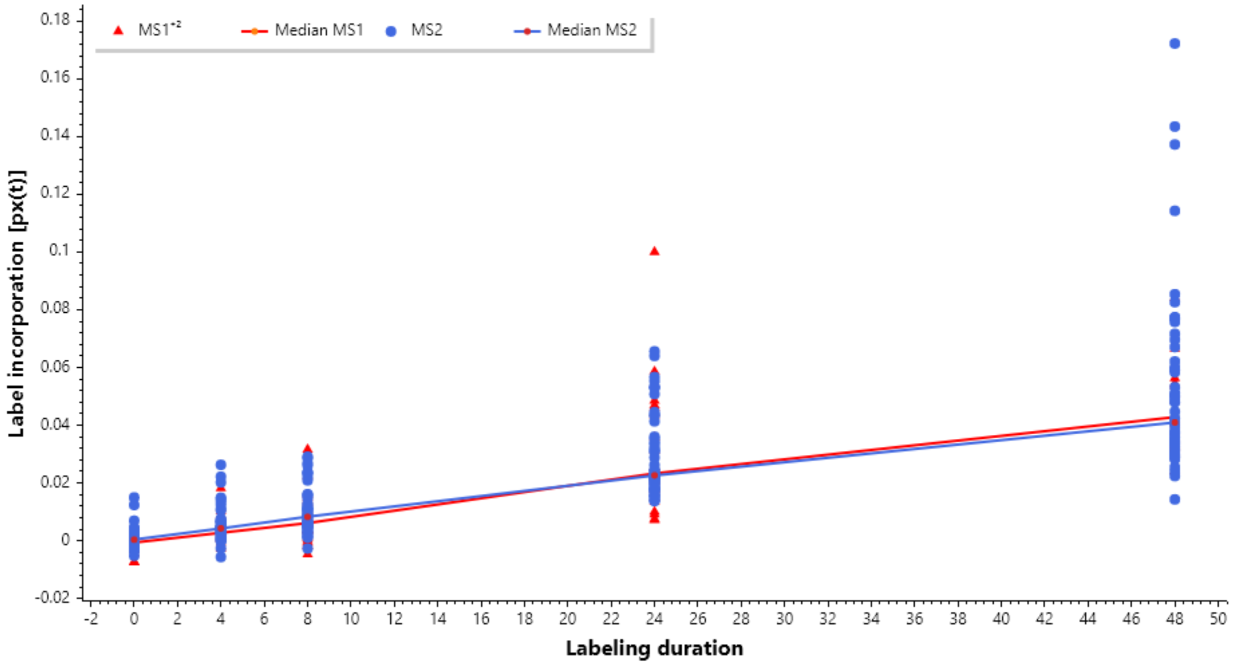


**Supplementary Figure 19**. The time course of label incorporation values, which are computed from precursor ions (MS1, represented by blue circles) and convolved fragment ions (MS2, represented by red triangles) over different labeling durations. The solid lines in red and blue show the median label incorporation values. For abundant peptides and precise intensity quantifications, the label incorporation values derived from MS1 and MS2 are comparable. Shown are the data for the peptide LDEAEQLALK$^{2+}$ of murine protein Myosin-8.

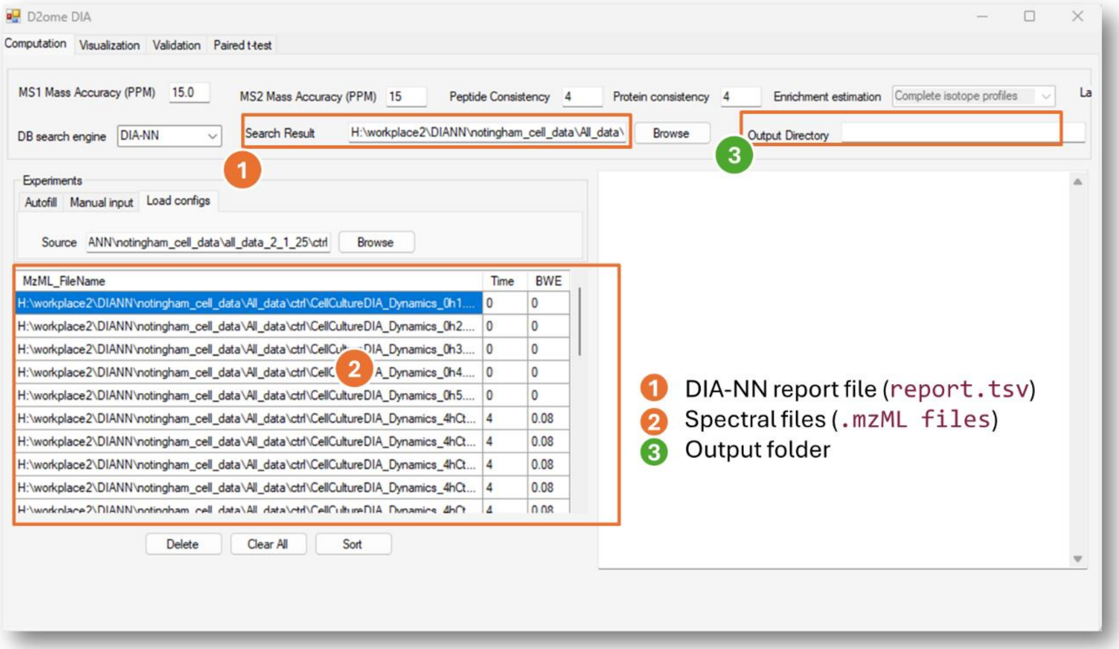


**Supplementary Figure 20.** The graphical user interface (GUI) to input data and parameters for the protein turnover rate estimations.

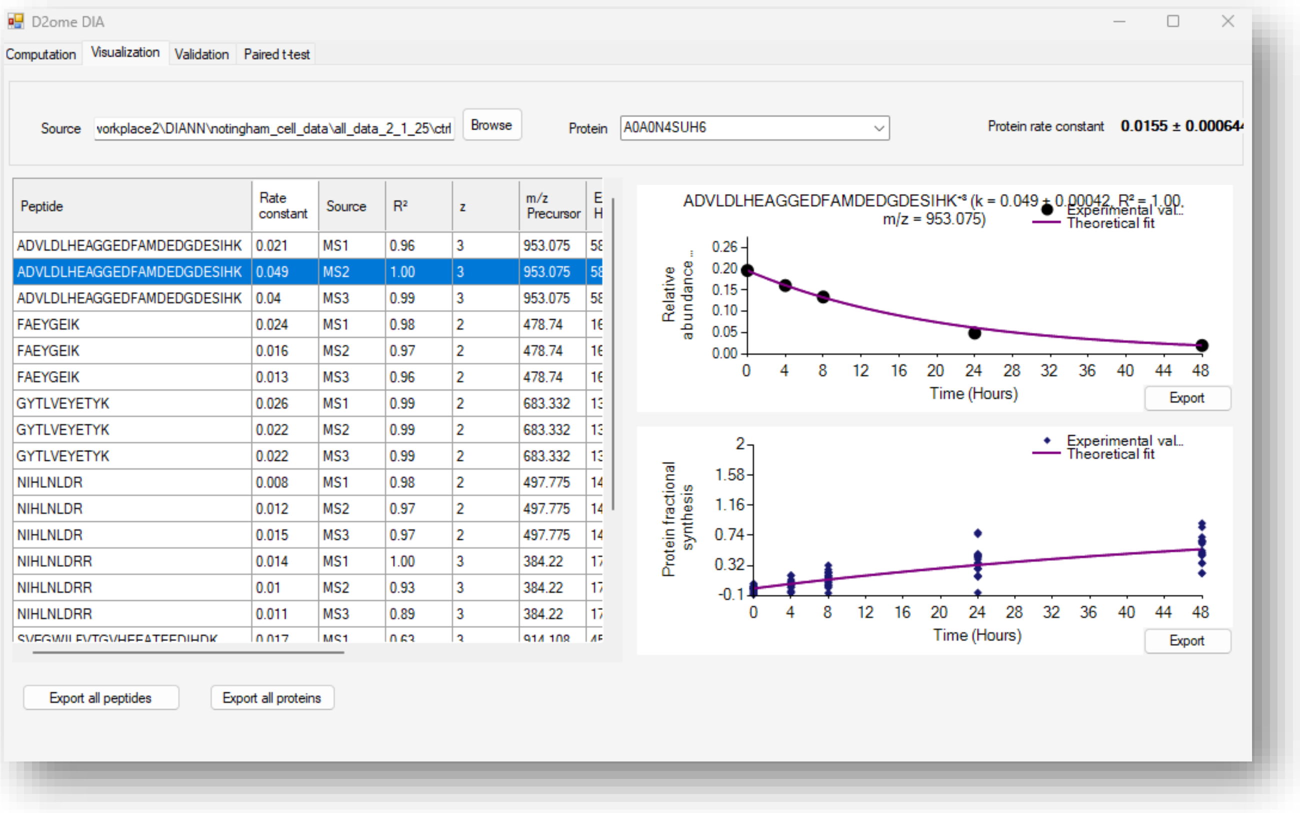


**Supplementary Figure 21.** The graphical user interface provides a detailed visualization of the results from protein turnover studies using metabolic $D_2O$-labeling and LC-MS experiments.

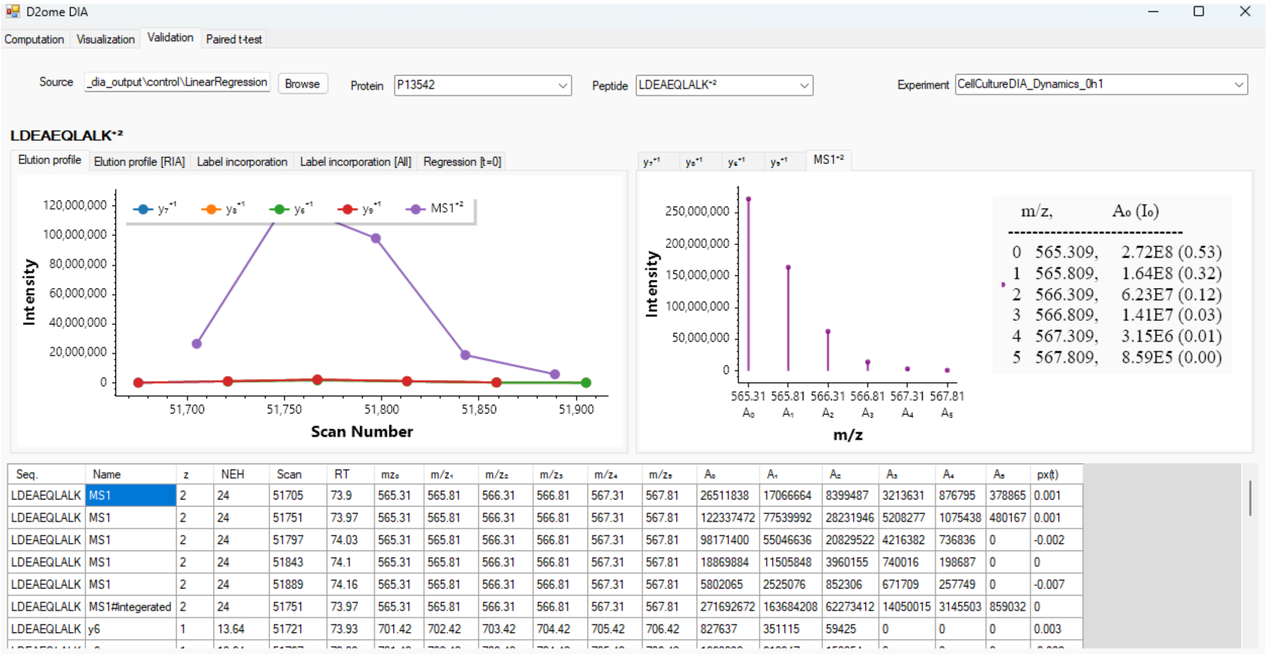


**Supplementary Figure 22.** The screenshot of the validation page of the d2ome-DIA graphical user interface. It shows the quantification (monoisotopic relative abundance, RA) results from MS1 and tandem mass spectra (MS/MS), absolute abundance of the monoisotopic peak, and retention times, mass-to-charge ratios of a peptide from liquid-chromatography mass spectrometry experiments at all labeling durations. A, B), C) D)XXXX

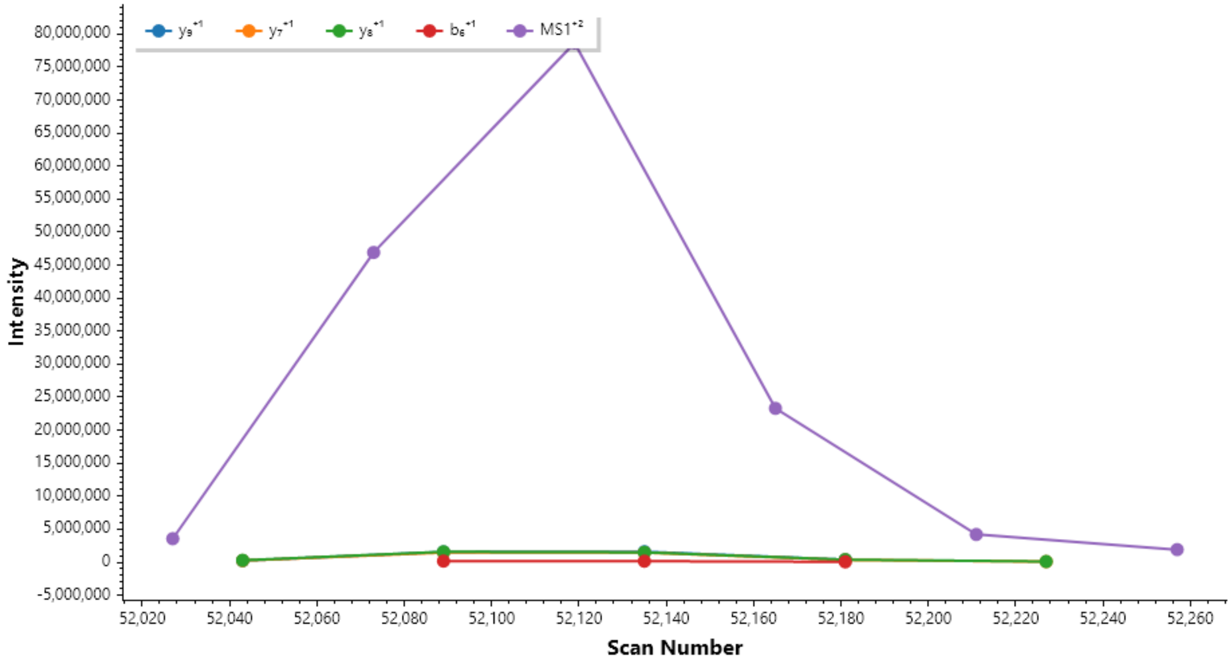


**Supplementary Figure 23.** Validation figures generated for the peptide LDEAEQLALK$^{2+}$ (MYH8_MOUSE) before the start of the labeling, derived from the "CellCultureDIA_Dynamics_0h1" experiment file. The scatter plot illustrates the raw abundance of the mono-isotope over the elution window of the peptide.

**Supplementary Table 1.** Database search engine report file format for D2ome-DIA input. The software accepts search results from any search engine, provided they are preprocessed as specified in the table below.

| Column name | Column description | Example |
|---|---|---:|
| File_Name | Experiment file name without the file extension | CellCultureDIA_Dynamics_4hCtrl4 |
| ProteinNames | Protein Names: UniProt accession(s) or protein identifiers | O88796 |
| ProteinGroups | Protein Group | O88796 |
| ModifiedPeptideSequence | Modified Peptide Sequence:Peptide sequence including post-translational or chemical modifications (UniMod notation) | (UniMod:1)AAFADLDLR |
| PeptideCharge | Charge state of the precursor peptide ion | 2 |
| PeptideMoverZ | Precursor peptide mass to charge ratio (m/z) | 517.2692871 |
| ElutionStartTime | Chromatographic elution start time | 112.1703262 |
| ElutionEndTime | Chromatographic elution end time | 112.4973297 |
| FragementIons | Observed fragment ions with annotation and m/z values | y6^1/702.3780518;y5^1/631.3409424;y4^1/516.3140259;... |
| FragementIonsScore | Correlation or confidence score(s) for the corresponding fragment ions | 0.8044459224;0.8886048794;0.9359041452;0.6546536684;0.6546536684;0; |
